# *N*-myristoyltransferase inhibition is synthetic lethal in MYC-deregulated cancers

**DOI:** 10.1101/2021.03.20.436222

**Authors:** Gregor A. Lueg, Monica Faronato, Andrii Gorelik, Andrea Goya Grocin, Eva Caamano-Gutierrez, Francesco Falciani, Roberto Solari, Robin Carr, Andrew S. Bell, Edward Bartlett, Jennie A. Hutton, Miriam Llorian-Sopena, Probir Chakravarty, Bernadette Brzezicha, Martin Janz, Mathew J. Garnett, Dinis P. Calado, Edward W. Tate

**Affiliations:** Department of Chemistry, Molecular Sciences Research Hub, White City Campus Wood Lane, Imperial College London, London W12 0BZ, UK; The Francis Crick Institute, 1 Midland Road, London NW1 1AT, UK; Institute of Integrative Biology, University of Liverpool, Liverpool, UK; Myricx Pharma Limited, Stevenage Bioscience Catalyst, Gunnels Wood Road, Stevenage, UK SG1 2FX; Experimental Pharmacology & Oncology Berlin-Buch, Robert-Rössle-Str. 10 13125 Berlin, Germany; Experimental and Clinical Research Center, Max Delbrück Center for Molecular Medicine and Charité – Universitätsmedizin Berlin, 13125 Berlin, Germany; Wellcome Sanger Institute, Cambridge, UK; Peter Gorer Department of Immunobiology, School of Immunology & Microbial Sciences, King’s College London, London, UK

## Abstract

Human *N*-myristoyltransferases (NMTs) catalyze N-terminal protein myristoylation, a modification regulating membrane trafficking and interactions of >100 proteins. NMT is a promising target in cancer, but a mechanistic rationale for targeted therapy remains poorly defined. Here, large-scale cancer cell line screens against a panel of NMT inhibitors (NMTi) were combined with systems-level analyses to reveal that NMTi is synthetic lethal with deregulated MYC. Synthetic lethality is mediated by post-transcriptional failure in mitochondrial respiratory complex I protein synthesis concurrent with loss of myristoylation and degradation of complex I assembly factor NDUFAF4, followed by mitochondrial dysfunction specifically in MYC-deregulated cancer cells. NMTi eliminated MYC-deregulated tumors in vivo without overt toxicity, providing a new paradigm in which targeting a constitutive co-translational protein modification is synthetically lethal in MYC-deregulated cancers.

**One-sentence summary:** *N*-myristoyltransferase inhibition leads to post-transcriptional complex I failure and cell death in MYC-deregulated cancers

## Introduction

*N*-myristoylation is a primarily co-translational and irreversible lipid modification at a protein N-terminal glycine, mediated in humans by the closely related enzymes *N*-myristoyltransferase (NMT) 1 and 2 (Fig. 1A) *(1)*. Myristoylation modulates membrane association *(2,3)*, protein stability *(4)* and interactions *(5)*, and proteomic and bioinformatic studies have identified over 100 substrates of NMT in the human proteome *(6–9)*. NMT has previously been proposed as a target in cancer *(10)*, but a rationale supporting a therapeutic index for NMT inhibition (NMTi) is yet to be identified. Many prior studies have been limited by availability of potent and selective NMT inhibitors *(11)*, and focused on individual NMT substrates *(12)* rather than addressing the system-wide consequences of NMTi across multiple cellular pathways, and its interactions with dynamic protein turnover. Recent discovery of the first potent and selective dual NMT1/NMT2 inhibitors provides a new avenue to address pharmacological validation of NMT as a target in cancer *(13)*, and we hypothesized that increased dependency on NMT could arise at the system level from deregulation of specific oncogenes. Here we combined large-scale screening of cancer cell lines against a panel of NMTi with systems-level analysis of cellular response to reveal that deregulation of MYC or MYCN renders cancer cells acutely sensitive to NMT inhibition. We find that NMT synthetic lethality in the context of MYC deregulation is mediated by rapid post-transcriptional failure in complex I protein synthesis, associated with NMTi-mediated depletion of myristoylated complex I assembly factor NDUFAF4, followed by mitochondrial dysfunction. NMTi eliminated patient-derived xenograft tumors in vivo without overt toxicity, providing a mechanistic framework for NMTi as a novel targeted cancer therapy. This new paradigm for targeting a constitutive protein modification in MYC deregulated cancer has potentially broad applications, since MYC remains undruggable despite being among the most commonly deregulated oncogenes *(14,15)*.

**Figure 1:**
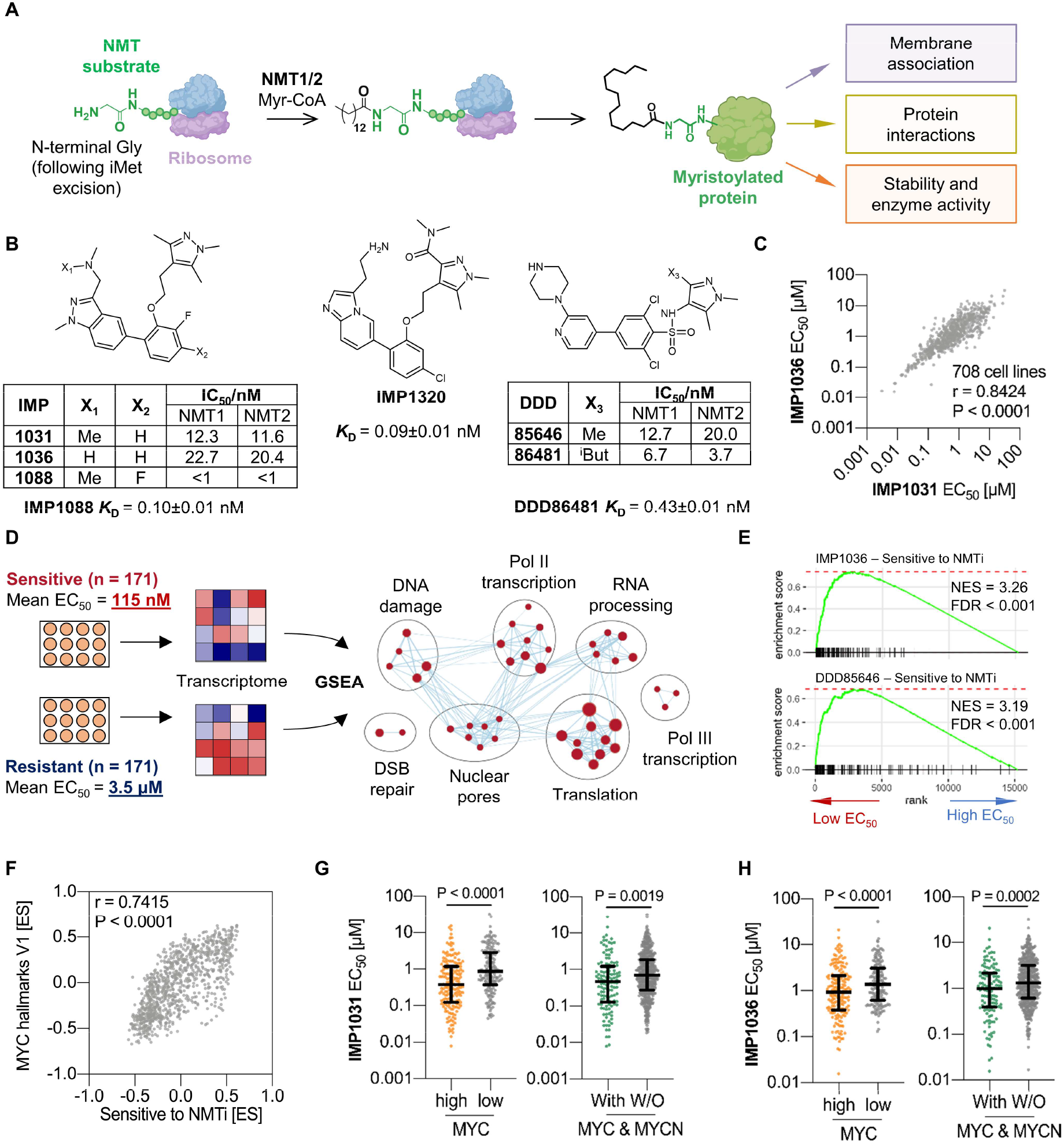
MYC deregulation and expression predict sensitivity to NMT inhibition. **(A)** Human NMT1 and NMT2 catalyze protein myristoylation of specific substrates during peptide elongation at the ribosome, leading to varied functions for the NMT substrate. **(B)** Chemical structure and inhibitory potency against human NMT1 and NMT2 for the NMT inhibitors used in this study. **(C)** Correlation of the EC_50_ values of IMP1031 and IMP1036 across 708 screened cancer cell lines (Spearman rank correlation). **(D)** Strategy to identify biological pathways enriched in sensitive cell lines, shown for IMP1031; GSEA was performed on COSMIC microarray data between sensitive and resistant quartiles. **(E)** The ‘Sensitive to NMTi’ gene set derived with IMP1031 was validated against independent cell line screen data for IMP1036 and DDD85646. **(F)** Correlation by GSVA between ‘Sensitive to NMTi’ and ‘MYC hallmark V1’ gene sets for COSMIC cell lines (Spearman rank correlation). **(G)** Left: EC_50_ values for cell lines screened against IMP1031, divided by quantiles into high and low expressers of MYC. Right: EC_50_ values grouped for cell lines with or without alterations in MYC and/or MYCN loci. **(H)** Analysis as for (G) using screen data for IMP1036 (Wilcoxon rank test; bars show median and IQR).

## Results

### MYC deregulation links to NMTi sensitivity

To identify susceptible cancer subtypes, we screened 708 cancer cell lines with extensive genomic and transcriptomic annotation *(16)* against three NMTi (IMP1031, IMP1036 and DDD85646; Fig. 1B) representative of two distinct NMTi chemotypes previously shown to possess excellent in-cell selectivity *(11)*. The remarkably wide (10^4^) range of susceptibility (IC_50_), excellent correlation between inhibitors and a proportional shift to lower IC_50_ across cell lines for NMTi with higher biochemical potency (Fig. 1C) supported robust on-target selectivity for NMT, which was reproduced in discrete viability assays (Supplementary Fig. 1). However, susceptibility was not significantly associated with cancer-functional-events (CFEs) *(16)*, mutations in known NMT substrates *(6,8,16)*, or expression of NMT1 or NMT2 (Supplementary Fig. 2), consistent with our hypothesis that sensitivity to NMTi has a non-trivial origin. Transcription of gene sets related to translation, RNA transcription and processing, DNA damage and repair, and nuclear import and export *(16)* were enriched in cell lines sensitive to IMP1031 (Gene Set Enrichment Analysis (GSEA) *(17)*, 171 cancer cell lines per group; Fig. 1D), and intersection of the leading-edge genes of the top 10 gene sets by normalized enrichment score (NES) (Supplementary Fig. 3) defined a consensus ‘Sensitive to NMTi’ gene set (Supplementary Table 1) robustly enriched in cell lines sensitive to IMP1036 or DDD85646, confirming the NMTi dependence of this transcriptional signature (Fig. 1E).

The ‘Sensitive to NMTi’ gene set is highly correlated with multiple gene signatures *(18)* upregulated by dysregulated MYC and anticorrelated with MYC downregulated gene signatures (Supplementary Fig. 4A) independent of genes associated with growth and proliferation, whilst other oncogenic signaling pathways (e.g. SRC, WNT/β-catenin) were not significantly associated (Supplementary Fig. 4B). Gene set variation analysis (GSVA) *(19)* confirmed a strong correlation between MYC Hallmark and ‘Sensitive to NMTi’ gene sets across the Catalogue Of Somatic Mutations In Cancer (COSMIC) cell lines, strongly implicating a MYC-driven transcriptional program in sensitivity to NMTi (Fig. 1F). Furthermore, MYC expression, or mutation, amplification, or chromosomal rearrangement in *MYC* and/or *MYCN* (a MYC paralogue commonly associated with neuroblastoma *(20)*) were predictive for NMTi sensitivity (Fig. 1G, H), and directly correlated with enriched expression of the ‘Sensitive to NMTi’ gene set (Supplementary Fig. 4C, D). Taken together, these analyses identified an unanticipated liability of MYC deregulated cancer cells to NMTi.

### NMTi is synthetic lethal in high-MYC cancers

To test the hypothesis that MYC deregulation is synthetic lethal with NMTi, we determined the sensitivity of P493-6 immortalized B cells carrying tetracycline-inducible MYC expression *(21)* to IMP1088 (Fig. 1B, Supplementary Fig. 5A), an NMTi with high potency against a panel of sensitive cancer cell lines (Supplementary Fig. 1) from the same class as IMP1031 and IMP1036 *(11,13)*. MYC was induced to a specific level (low, medium or high) over 24 hours *(21)* (Fig. 2A), and cells exposed to 100 nM IMP1088, sufficient to fully inhibit cellular NMT activity *(11)* (Supplementary Fig. 6), for 72 hours. High-MYC cells underwent a precipitous decrease in viability (Fig. 2B; Supplementary Fig. 7A), and >15-fold decrease in cell number (Fig. 2C) relative to DMSO-treated controls, whilst the impact of NMTi on medium-MYC cells was modest, and low MYC cells were largely unaffected by NMTi over 72 hours. Similar synthetic lethality was observed in neuroblastoma (NB) cell line MYCN-ER-SHEP, in which 4-hydroxytamoxifen (tam)-induced MYCN mimics the highly aggressive *MYCN* amplified form of clinical NB *(22)* (Fig. 2D-F). NMTi strongly induced apoptosis within 48 hours (Supplementary Fig. 7B) and a subsequent rapid decrease in cell viability by 72 hours (Fig. 2F) exclusively in the presence of elevated MYCN expression. Each of these outcomes was recapitulated with a chemically distinct, potent and selective NMTi (DDD86481 *(23)*, Fig. 1B; Supplementary Fig. 5B, 6 and 8), confirming the role of NMT inhibition in synthetic lethality.

**Figure 2:**
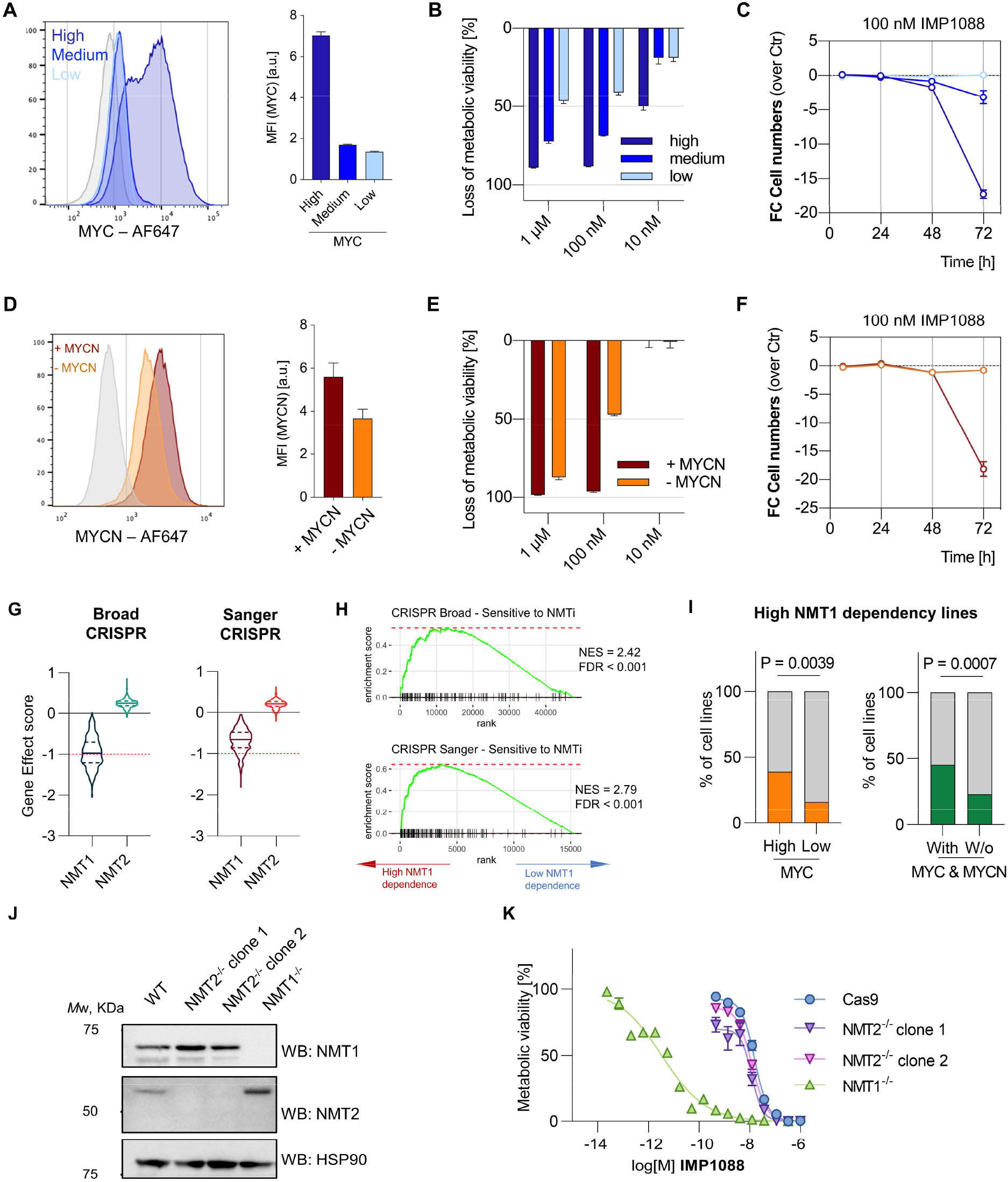
NMTi is synthetically lethal with MYC or MYCN induction and driven by NMT1 inhibition. **(A)** Flow cytometry analysis and quantification of MYC induction in P493-6 cells. **(B)** Dependence of IMP1088 (100 nM) toxicity on MYC in P493-6 cells (CellTiter-Blue assay). **(C)** Fold-change in cell numbers in MYC induced P493-6 with 100 nM IMP1088, measured over time by flow cytometry. **(D)** Flow cytometry analysis and quantification of MYCN induction in SHEP cells. **(E)** Dependence of IMP1088 (100 nM) toxicity on MYC in SHEP cells (CellTiter-Blue assay). **(F)** Fold-change in cell number (100 nM IMP1088 vs control) with or without MYCN induction in SHEP cells measured over time by flow cytometry. **(G)** Gene effect scores for NMT1 and NMT2 for sgRNA libraries, analyzed using the Broad pipeline. Red dashed line: median gene effect scores of genes classified as core essential. **(H)** Enrichment of the ‘Sensitive to NMTi’ gene set in NMT1-dependent cancer cell lines (negative versus positive gene effect scores by quantiles, Broad DepMap) or cell lines designated dependent on NMT1 (Sanger Project Score). **(I)** Representation of NMT1-dependent cell lines among lines expressing high MYC (by quantiles), or with structural alterations in *MYC* or *MYCN* loci (Fisher-Exact test, Sanger Project Score). **(J)** Western blot for NMT1 or NMT2 in CRISPR-Cas9 mediated knockout and wild type HeLa cells. **(K)** Sensitization to NMT inhibition by IMP1088 in HeLa NMT1^-/-^ and NMT2^-/-^ cells (CellTiter-Blue assay). Data in panels A-F and K are shown as mean ± s.e.m. of n = 3 biological replicates.

To investigate if MYC deregulation represents a pan-cancer vulnerability to NMTi, we performed a screen using clonogenic analysis of 3D-cultured Patient-Derived (PD) cells spanning 18 cancer types across 50 patients using the potent NMT inhibitor IMP1320 (Fig. 1B, Supplementary Fig. 5C, and 9). Functional analyses on genes positively and negatively correlated with IMP1320 IC_50_ (Supplementary Fig. 10) revealed biologically relevant functional pathways. We found that low expressed genes in highly sensitive PD cancer cells enriched for functional terms linked to cell adhesion and membrane components, whereas highly expressed genes enriched for functional terms related to RNA and DNA processes (Supplementary Fig. 10A). Ingenuity Pathway Analysis (IPA) revealed MYC as the highest scoring gene in the most sensitive PD lines (Supplementary Fig. 10B) and that MYC and MYCN target gene expression correlated to NMTi sensitivity (Supplementary Fig. 10C). Consistent with an overactivated MYC status in highly NMTi sensitive PD cancer cells, let-7, a miRNA inhibiting MYC expression *(24)*, was active in less sensitive PD cancer cell lines. Collectively, these data support MYC-deregulation as a pan-cancer vulnerability to NMTi.

### Deregulated MYC drives NMT1 dependence

All potent human NMT inhibitors reported to date are dual NMT1 and NMT2 inhibitors due to the high homology of these isoforms in the catalytic domain *(9,25)*. Whole genome CRIPSR knockout screens encompassing >500 cancer cell lines (Cancer Dependency Map, DepMap *(26,27)*) processed with a common pipeline *(28)* revealed that NMT1 is required for optimal proliferation of most cancer cell lines, whereas NMT2 appears to lack essentiality in any cell line (Fig. 2G). Cancer cell lines with greater dependence on NMT1 expression are enriched in the ‘Sensitive to NMTi’ gene set (Fig. 2H) and are also overrepresented in cell lines classified by high MYC expression and/or structural alterations in MYC or MYCN (Fig. 2I, Supplementary Fig. 11). To explore this predominant dependence on NMT1 we generated *NMT1* or *NMT2* homozygous knockouts using CRISPR-Cas9 in HeLa cells (Fig. 2J, Supplementary Fig. 12). NMT1 knockout conferred 1000-fold greater sensitivity to IMP1088 (shifting the EC_50_ value from 10 nM to 10 pM) whereas NMT2 knockout had no impact (Fig. 2K), confirming that NMT2 expression has negligible impact on NMTi sensitivity.

### NMTi impacts MYC-driven proteome dynamics

*N*-myristoylation is predominantly co-translational *(6)* and irreversible *(29)*, and we hypothesized that synthetic lethality with MYC deregulation results from the interplay between the cellular state induced by NMTi and the profound influence of MYC on protein homeostasis. MYC upregulation drastically increases ribosomal and mitochondrial biogenesis *(30)* and induces a strong proteotoxic stress response *(31–33)*, but its quantitative impact on proteome dynamics has not been reported to date. We applied a triplex SILAC strategy *(34)* to determine changes in protein half-life *(35)* and rates of protein synthesis and degradation *(36)* in high-versus medium-MYC P493-6 cells in the presence or absence of 100 nM IMP1088 over 4, 8, 16 and 24 hours (Fig. 3A; Supplementary Fig. 13). The overall bias of protein half-lives was significantly shorter in high-MYC vs medium-MYC cells (Fig. 3B; Supplementary Fig. 14A,B) but despite a modest overall trend toward reduced NMT substrate half-life in high-versus medium-MYC cells (Fig. 3C) it was notable that high MYC expression did not uniformly accelerate protein turnover (Supplementary Fig. 14C). 2D enrichment *(37)* analysis revealed concordant changes in protein synthesis rate and mRNA expression by RNA-seq (Fig. 3D), with a strong bias toward translation, mitochondrial biogenesis and RNA metabolism in high-MYC relative to medium-MYC cells (Fig. 3D).

**Figure 3:**
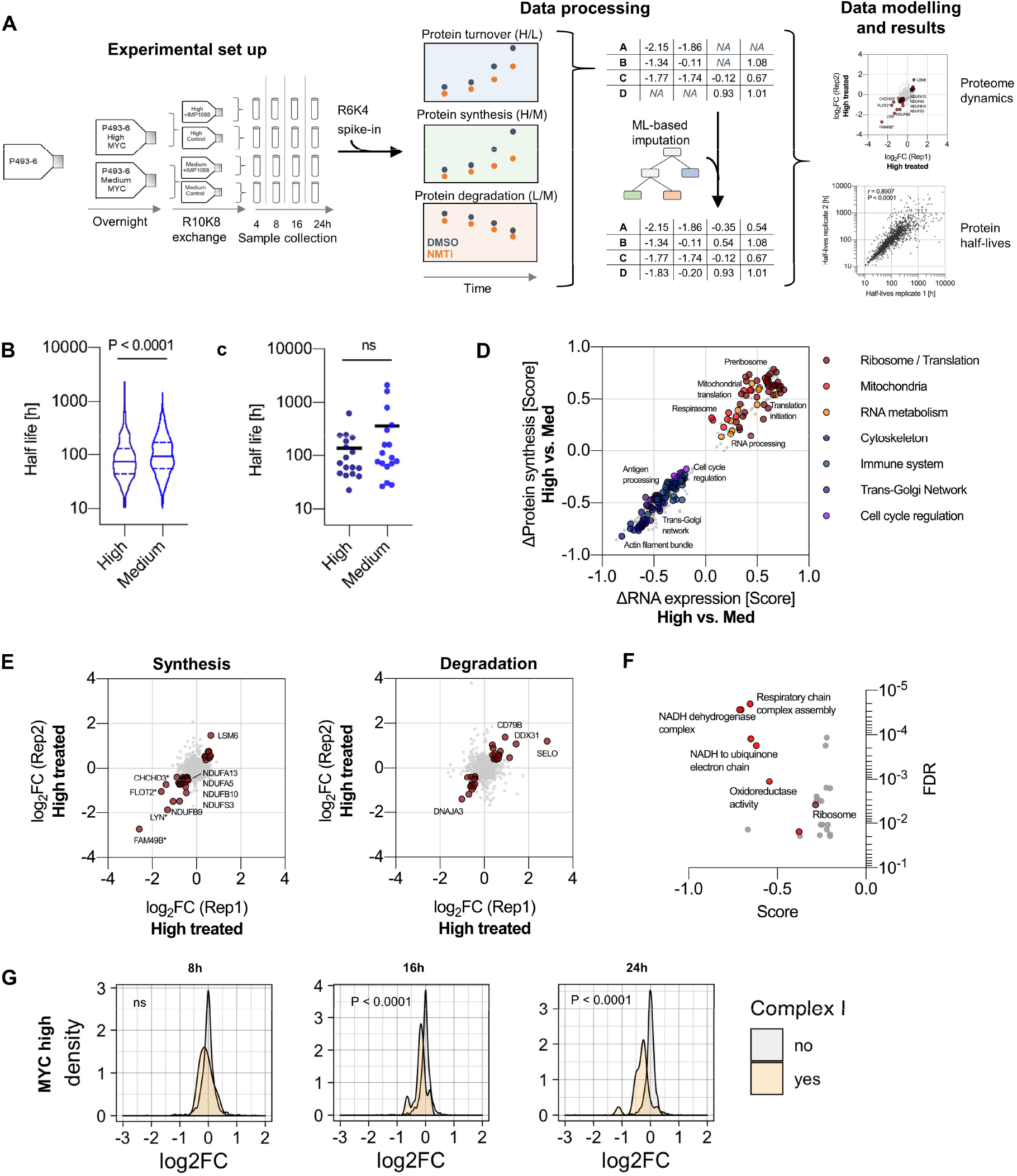
NMT inhibition impacts mitochondrial complex I protein dynamics in MYC deregulated cells. **(A)** P493-6 cells at specified levels of MYC induction were transferred into heavy (R10K8) media containing 100 nM IMP1088, or DMSO control and incubated for 4, 8, 16 or 24 h. Medium (R6K4) labeled combined proteome was spiked into all samples prior to analysis to enable relative quantification across samples, and data processed as described in the methods section. **(B)** Half-lives of proteins identified in both high-and medium-MYC cells (n = 592, Wilcoxon rank test). **(C)** Half-lives of NMT substrates identified in both high-and medium-MYC cells (n = 17, Wilcoxon rank test). **(D)** 2D-enrichment between changes in mRNA abundance and protein synthesis in high-MYC versus medium-MYC cells; FDR threshold ≤ 0.1%. **(E)** Effect of IMP1088 on rates of protein synthesis (H/M SILAC ratio) and degradation of pre-existing proteins (M/L SILAC ratio) in high-MYC cells (asterisk indicates NMT substrate). **(F)** 1D-enrichment on changes in synthesis rate, showing impact on complex I of the mitochondrial respiration chain in high-MYC cells; FDR threshold ≤ 2%. **(G)** Effect of IMP1088 (100 nM) on the synthesis of proteins of mitochondrial complex I over time in high-MYC cells (Wilcoxon rank test).

We next investigated the impact of NMTi (100 nM IMP1088) *(36)* (Supplementary Fig. 14D,E and Supplementary Methods) and observed that synthesis and degradation rates of a subset of proteins were strongly impacted by NMT inhibition in high-MYC cells (Fig. 3E), with 1D-enrichment on differential synthesis suggesting a particularly strong impact on the NADH dehydrogenase complex, also known as mitochondrial complex I (Fig. 3F). NMTi induced a time-dependent decrease in the rate of synthesis of proteins of complex I in high-MYC cells (Fig. 3G), and whilst similar overall trends were seen in medium-MYC cells (Supplementary Fig. 15) high-MYC cells displayed a much higher baseline demand for mitochondrial protein synthesis (Fig. 3D). Furthermore, 2D enrichment analysis suggested that NMTi caused post-transcriptional failure in complex I protein synthesis, without corresponding downregulation of mRNA (Supplementary Fig. 15D).

### NMTi drives mitochondrial dysfunction in high MYC cancer cells

Since MYC expression drives increased mitochondrial biogenesis, we hypothesized that failure to synthesize complex I upon NMTi causes a catastrophic failure in mitochondrial function in high-MYC cancer cells *(38)*, leading to cell death. Mitochondrial potential was significantly decreased, and superoxide generation increased by IMP1088 treatment (100 nM, 18 hours) only in high-MYC P493-6 cells (Supplementary Fig. 16A), prompting us to measure the impact of NMTi on mitochondrial respiration. As previously reported *(39,40)*, high MYC expression increases mitochondrial respiration (Supplementary Fig. 16B). In line with progressive loss of complex I synthesis (Fig. 3G), NMTi induced a time-dependent reduction in basal and maximal respiration, ATP production and spare respiratory capacity in high-MYC cells with effects already observable after 12 h treatment, whilst medium-MYC cells were unaffected (Fig. 4A,B; Supplementary Fig. 16C,D). MYC-dependent mitochondrial phenotypes were replicated with DDD86481 (Fig. 4A,B), further confirming the effect of NMTi. Notably, IMP1088 and DDD86481 induced similar impacts on mitochondrial function in PD LY11212 diffuse large B-cell lymphoma (DLBCL) cancer cells *(41)* (Fig. 4C,D; Supplementary Fig. 16E, 17). These cancer cells are derived from a patient with multi-chemotherapy resistant lymphoma carrying *MYC* and *BCL2* translocations, characteristic of so-called “double hit” lymphomas which have the least favorable clinical outcomes among all DLBCL *(42)*.

**Figure 4:**
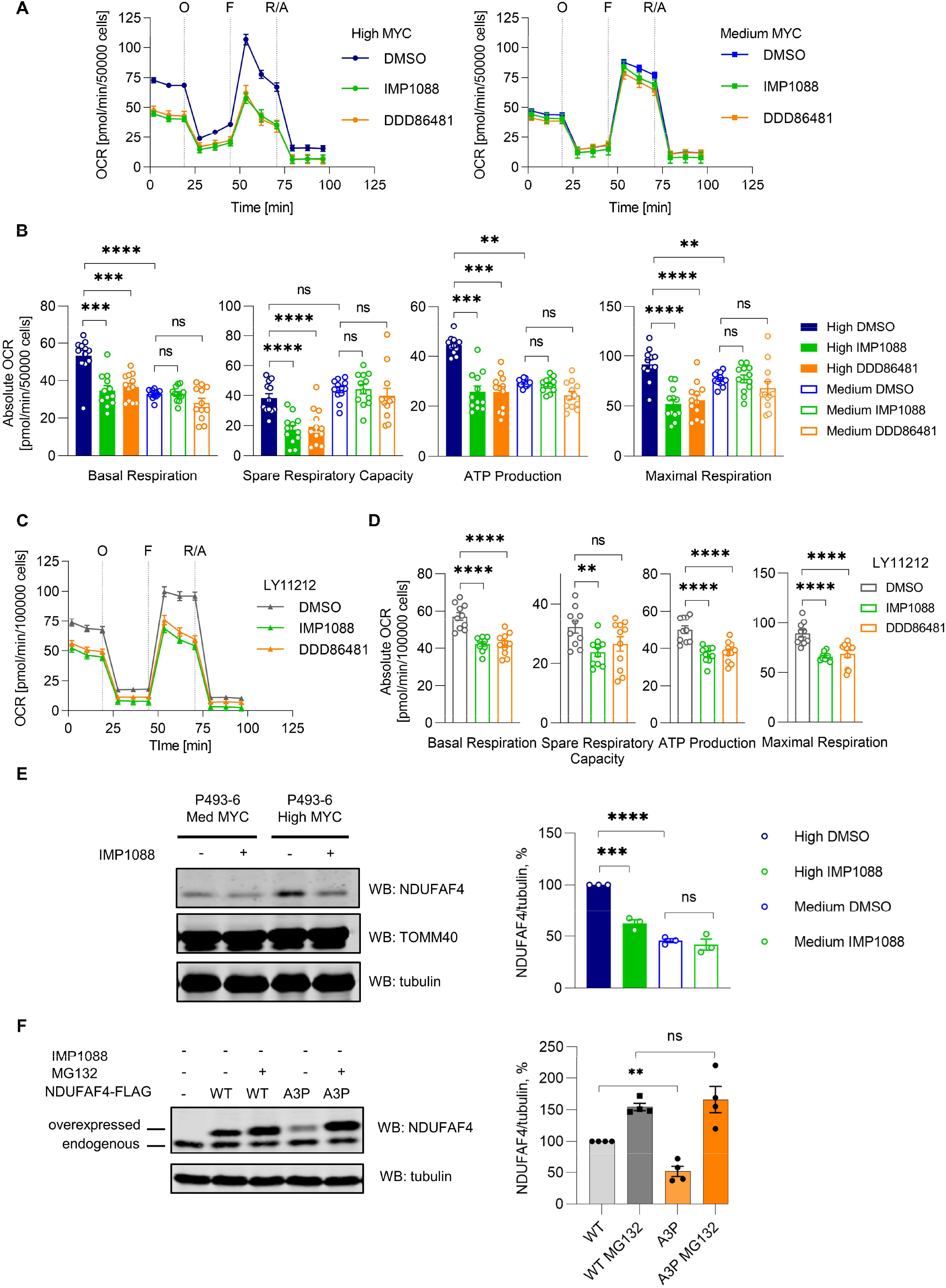
NMT inhibition severely impairs mitochondrial respiration in high-MYC cells. **(A)** Oxygen consumption rate (OCR) of P493-6 cells expressing medium and high MYC levels upon treatment with IMP1088 (100 nM) or DDD86481 (1 µM) for 18 h (measured using Seahorse XFe96 analyzer), vehicle control was DMSO. O: oligomycin, F: FCCP, R/A: rotenone and antimycin A. **(B)** Parameters of mitochondrial function in P493-6 cells calculated using data from (A). **(C)** OCR of LY11212 PD cancer cells upon treatment with IMP1088 (100 nM) or DDD86481 (1 µM) for 18 h. **(D)** Parameters of mitochondrial function in LY11212 PD cancer cells calculated using data from (C). **(E)** Western blot analysis of NDUFAF4 levels in P493-6 cells (medium or high MYC) with and without IMP1088 treatment (100 nM, 18 h). Tubulin was used as a loading control. For quantification, normalization was performed by dividing the NDUFAF4 antibody signal by the tubulin antibody signal. **(F)** Western blot analysis of C-terminally FLAG-tagged NDUFAF4 (WT and A3P mutant) expressed in HEK293 cells. MG132 (10 µM) was used to inhibit the proteasome. Tubulin was used as a loading control. For quantification, normalization was performed by dividing the NDUFAF4 antibody signal by the tubulin antibody signal. Data in **A-F** are shown as mean ± s.e.m. of n = 3 biological replicates. ns: not statistically significant, * *P*<0.05, ** *P*<0.01, *** *P*<0.001, **** *P*<0.0001 (two-way ANOVA – panels **A-D**, t-test: two-tailed, unpaired – panels **E, F**).

NDUFAF4 is a mitochondrial NMT substrate that functions as a complex I assembly factor and loss of NUDFAF4 is associated with impaired complex I expression *(6,43)*. We previously identified NDUFAF4 as a human NMT substrate *(6)*, and subsequent studies have shown that non-myristoylated NDUFAF4 is subject to degradation via the glycine N-degron pathway *(4)*. We found that NDUFAF4 protein levels were specifically and significantly reduced upon NMTi in high-MYC cells whereas another known mitochondrial NMT substrate TOMM40 was unaffected (Fig. 4E). Recently, it was reported that patients carrying a single Ala3Pro mutation in NDUFAF4 suffer a specific mitochondrial complex I assembly defect leading to onset of Leigh syndrome *(44)*. We hypothesized that this mutation adjacent to the Gly2 *N*-myristoylation site abolishes NDUFAF4 *N*-myristoylation, leading to NDUFAF4 degradation through the glycine N-degron pathway. To test this hypothesis, we expressed wild type NDUFAF4 or NDUFAF4[Ala3Pro] with a C-terminal FLAG tag in human embryonic kidney (HEK293) cells, and found that NDUFAF4[Ala3Pro] expression was significantly reduced relative to wild type, which could be rescued by proteasome inhibition (Fig. 4F). NDUFAF4[Ala3Pro] N-terminal peptide is not a substrate for recombinant human NMT in contrast to efficient myristoylation of wild type NDUFAF4 peptide (Supplementary Fig. 18), and NDUFAF4[Ala3Pro] protein was not metabolically labelled by myristate analogue YnMyr *(6)* in cells (Supplementary Fig. 19). These data support the hypothesis that impaired NDUFAF4 myristoylation upon NMTi leads to complex I assembly defects in high MYC cells, and furthermore suggest that failure to myristoylate NDUFAF4 is on its own sufficient to impair physiological complex I assembly in Leigh syndrome patients.

### NMTi suppresses *MYC* deregulated tumors

We next examined the impact of NMTi in PD double hit high-grade B-cell lymphomas LY11212 *in vivo*, which were exquisitely sensitive to both IMP1088 (EC_50_ 5 nM) and DDD86481 (EC_50_ 16 nM) *in vitro* (Supplementary Fig. 20). LY11212 cells were engrafted subcutaneously into NOD scid gamma (IL2R-NSG) mice and tumors established over three days; mice were then treated with vehicle or DDD86481 at 25 mg/kg once per day for up to 11 days by intraperitoneal (IP) injection (10 mice per group). DDD86481 was selected for *in vivo* experiments since its pharmacokinetic profile predicted exposure above EC_50_ for the majority of dosing period (Supplementary Fig. 21). NMTi treatment resulted in profound inhibition of tumor growth in all cases, with the majority of NMTi treated animals showing no palpable tumor by the end of the experiment, whereas tumors grew in all vehicle-treated controls reaching humane endpoint within 13 days (Fig. 5A). Furthermore, the dose of NMTi was well tolerated by the animals, with no animal exhibiting overt signs of toxicity or weight loss throughout the experiment (Fig. 5B). We further profiled orally bioavailable NMTi IMP1320, engrafting double-hit DLBCL DoHH2 cancer cells into IL2R-NSG mice to establish tumors to a volume of 100-150 mm^3^, after which mice were treated with IMP1320 at 50 mg/kg or vehicle in two different groups once per day for 10 days by oral gavage (10 mice per group) and tumor volumes measured twice-weekly. NMTi treatment resulted in elimination of tumor in all animals, with no palpable tumors by day 22 of the experiment, while tumors grew in all vehicle-treated controls (Fig.5C). No substantial effect on body weight was observed, suggesting this dose is well-tolerated (Fig. 5D).

**Figure 5:**
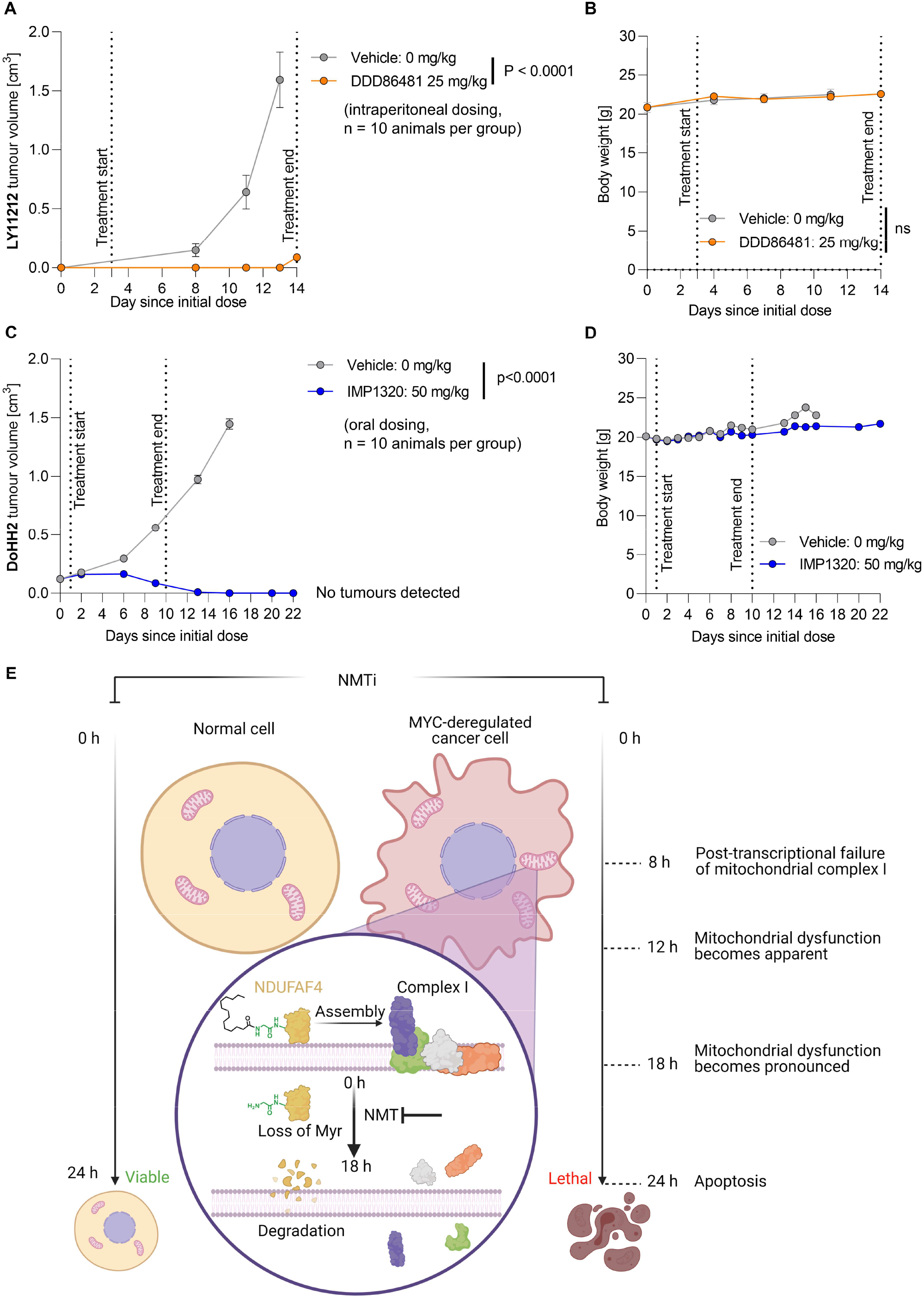
NMT inhibition abolishes tumor growth in vivo. **(A)** Impact of intraperitoneally administered DDD86481 on the growth of PDX in vivo using PD cancer cells LY11212 that carry a *MYC* translocated allele (n = 10 mice per group; error bars represent mean ± s.e.m.; time-adjusted ANOVA). **(B)** Change in mouse body weight between the start and end point of the experiments for the LY11212 PDX, comparing treated and control (t-test, paired). **(C)** Impact of orally administered IMP1320 on the growth of DoHH2 cancer cells in vivo (n = 10 mice per group; error bars represent mean ± s.e.m.; time-adjusted ANOVA). **(D)** Change in mouse body weight between the start and end point of the experiments for the DoHH2 xenograft, comparing treated and control. Error bars represent mean ± s.e.m. **(E)** Proposed mode of action of NMT inhibition in MYC-deregulated cancer cells and its impact on mitochondrial function.

## Conclusions

The remarkable and unexpected synthetic lethality of NMT inhibition in MYC deregulated cancers offers a new paradigm for targeting a constitutive co-translational modification in cancer, in which the cellular state induced by NMTi interacts with abnormal proteome dynamics of the MYC oncogenic program to provide a useful therapeutic window *(45)*. Our proteome dynamics data are consistent with previous observations in *Drosophila* connecting loss of genes involved in protein biogenesis with increased translation stress *(46)*, although the synthetic lethality observed for NMTi is both more selective and more potent than for inhibitors of previously reported protein synthesis-related targets in high-MYC cells *(47,48)*. Our data further suggest that inhibition of NMT1 (the paralogue most similar to the single NMT in lower eukaryotes) primarily drives MYC synthetic lethality, and that NMT2 expression plays little role in NMTi sensitivity in general. Indeed, regardless of its expression in many tumors, NMT2 appears to be neither an oncogene nor a tumor suppressor, and a clear role for this paralogue remains to be determined.

A recent report has proposed that the cytotoxicity of NMTi in B cell lymphomas arises from impaired B cell receptor (BCR) signaling through Src-family kinase (SFK) SRC and LYN degradation, both known NMT substrates *(12)*. We tested this hypothesis through gene-effect score correlation analysis between SFK and NMT1 gene knockout across cancer cell lines or specifically in leukemias and lymphomas (Supplementary Fig. 22) but found no correlations. Furthermore, we found that components of the BCR signaling pathway were downregulated only from the point where cell death is initiated (24 h after NMTi treatment, Supplementary Fig. 23). We conclude that the mechanism of NMTi synthetic lethality is largely independent of SFK/BCR signaling, particularly when compared to the profound and early impact of NMTi on proteome dynamics and mitochondrial complex I in MYC-deregulated cells.

Mitochondrial dysfunction is both a hallmark and a liability of MYC deregulation, and the impact of NMTi on mitochondria is remarkably rapid in high-MYC cells, depleting complex I protein production within 8 hours and mitochondrial respiration by 12 hours, preceding initiation of cell death by at least 12 hours. This is faster than the half-life of the large majority of proteins and NMT substrates measured in high-MYC cells, suggesting that complex I synthesis is highly sensitive to loss of *N*-myristoylation on newly synthesized proteins. The mechanism by which this precipitous decrease in complex I synthesis rate is decoupled from mRNA expression, and the extent to which dysregulation of NMT substrates with roles in mitochondria *(1)* or suppression of oxidative stress *(2)* impact mitochondrial sufficiency are interesting questions for future studies. Although the mechanisms by which NMTi induces cancer cell death operate at the level of the system as a whole, loss of myristoylation of complex I assembly factor NDUFAF4 alone is sufficient to drive physiological complex I defects in humans, as seen in Leigh syndrome (Fig. 5E).

We note that alterations in the proximal MYC network *(49)* correlate not only with worse clinical outcome *(50)* but also enrichment of both the MYC Hallmark gene set and the NMTi sensitivity signature (Supplementary Fig. 24). Correlation between these two signatures across TCGA cohorts (Supplementary Fig. 25) supports the potential to target NMT in a clinical setting. The advent of potent and selective human NMT inhibitors has proven essential to facilitate robust screening and system-level studies and to establish novel markers for NMTi sensitivity in cancer. Whilst it is clear that NMTi will not be without risk of toxicity in patients, our data suggest that a significant therapeutic window exists to target MYC deregulated cancers with NMTi. We expect that future refinement of dose schedules and understanding of dose-limiting toxicity will enable clinical development of NMT inhibitors targeting high-MYC cancers.

## Author contributions

EWT and DPC conceived the study. GAL developed and applied the data analysis pipelines for NMTi sensitivity, gene essentiality, SILAC, TCGA, performed RNA-seq experiments with help from MLS and PC, and metabolic viability and FACS experiments. MF performed metabolic viability and FACS experiments for the SHEP cell line with NMTi and for PDX with DDD86841, generated CRISPR-Cas9 knockout of NMT1/2 in HeLa cells, prepared SILAC samples, performed mitochondrial potential and ROS assays and carried out *in vivo* mouse experiments with The Francis Crick Institute Biological Research Facility. AG performed in vitro and cellular NMTi inhibition assays, YnMyr profiling by in-gel fluorescence, Seahorse mitochondrial analysis, NDUFAF4 cloning, mutagenesis, expression and biochemical experiments, and contributed to manuscript writing. AGG assisted with proteomics data analysis and data deposition. EC-G and FF performed bioinformatics analysis on the patient-derived cell screen. ASB designed IMP1320. JAH designed and synthesized IMP1031 and IMP1036. MLS and PC provided support for bioinformatic analyses. RS and RC provided data on clonogenic analysis of 3D-cultured patient-derived cells and DoHH2 tumor experiments. BB and MJ generated and provided B cell lymphoma PDX lines. EB purified NMT1 and NMT2 and performed SPR experiments. MJG supervised collection and analysis of cell line screen inhibition data. EWT and DPC supervised the study and secured funding. EWT wrote the manuscript, with input from all authors.

## Acknowledgments

We thank Brigitte Wollert-Wulf (MDC, Berlin) for excellent technical assistance and Jens Hoffman (EPO, Berlin) for support in obtaining B cell lymphoma PDX. We thank Chi Van Dang (Ludwig Institute for Cancer Research, New York) for providing the P493-6 cell line and Michael D. Hogarty (University of Pennsylvania, Philadelphia) for providing the SHEP cell line. We thank Josephine Walton and Wouter Kallemeijn (Imperial College London) for support with data collation and presentation. We thank Cancer Research UK (C29637/A21451 and C29637/A20183 to EWT), Imperial College London (studentship award to GAL), the MRC (career development award MR/J008060/1 to DPC), the Deutsche Forschungsgemeinschaft (grant JA 1847/2-1 to MJ), and the Wellcome Trust (206194 to MJG). The Francis Crick Institute receives its core funding from Cancer Research UK (FC001057), the UK Medical Research Council (MRC; FC001057), and the Wellcome Trust (FC001057) (EWT and DPC). Figures 1A and 5E were created with biorender.com.

## Data availability

RNA-seq datasets have been deposited onto GEO with the dataset identifier GSE154336. The mass spectrometry proteomics data have been deposited to the ProteomeXchange Consortium via the PRIDE *(51)* partner repository with the dataset identifier PXD020318. All other data are available upon reasonable request to the corresponding authors.

### Code availability

An R script for the proteomics imputations and example from the experiment are provided for reproduction of the imputation.

