## Supplementary Code for "*N*-myristoyltransferase inhibition is synthetic lethal in MYC-deregulated cancers": ML_based_imputation.R

```
#The following R script allows for the reproduction of the ML-based multiple imputation for the
#proteomics dynamics experiment. An example data frame is provided and the following script allows
#to test different strategies of multiple imputation (by splitting the original data frame into lists of
#data frames that do not contain all of the original data) and different ML-based learners to impute the
#NAs in the data. The script was written (particularly in terms of column indices) for the experiment in the
#publication. However, it can be easily adopted for other types of data.

# Check if libraries for this script are installed ------------------------

#This section will verify if the R libraries (all available from CRAN) are installed - if not, it will install them.
if("tidyverse" %in% rownames(installed.packages()) == FALSE) {install.packages("tidyverse")}
if("magrittr" %in% rownames(installed.packages()) == FALSE) {install.packages("magrittr")}
if("foreach" %in% rownames(installed.packages()) == FALSE) {install.packages("foreach")}
if("mlr" %in% rownames(installed.packages()) == FALSE) {install.packages("mlr",
                                                                         dependencies = TRUE)}
if("gbm" %in% rownames(installed.packages()) == FALSE) {install.packages("gbm")}
 
# Libraries ---------------------------------------------------------------

#Loads in the libraries
library(tidyverse)
library(magrittr)
library(foreach)
library(mlr)
library(ParamHelpers)
library(gbm)

# Functions ---------------------------------------------------------------

#Functions that will be subsequently used

#Function will create a list of data frames, splitted into several subdataframes.
#Splits can be defined (default = 10), and if the rows should be shuffled prior splitting (default = TRUE).
CV_df_list_creater = function(df,
                              splits = 10,
                              randomise_rows = TRUE) {
  #if randomise_rows = TRUE the rows will be shuffled
  if (randomise_rows) {
    df_impute = df[sample(nrow(df)),]  
  } else {
    df_impute = df
  }
  #the data frame is split into two separate DFs with NAs and without NAs
  df_impute_w_NA = df_impute[!complete.cases(df_impute),]
  df_impute_wo_NA = df_impute[complete.cases(df_impute),]
  #two folds are defined based on pre-defined splits
  folds_w_NA = cut(seq(1, 
                       nrow(df_impute_w_NA)),
                   breaks = splits,
                   labels = FALSE)
  folds_wo_NA = cut(seq(1,
                        nrow(df_impute_wo_NA)),
                    breaks = splits,
                    labels = FALSE)
  #a list of dataframes is created
  df_list = foreach (i = 1:splits) %do% {
    #define the indices
    testIndex_wo_NA = which(folds_wo_NA == i,
                            arr.ind = TRUE)
    testIndex_w_NA = which(folds_w_NA == i, 
                           arr.ind = TRUE)
    #make the two data sets
    sampling_wo_NA = df_impute_wo_NA[-testIndex_wo_NA, ]
    sampling_NA = df_impute_w_NA[-testIndex_w_NA, ]
    #compine the two data frames together
    sampling = rbind(sampling_wo_NA,
                     sampling_NA)
    return(sampling)
  }
}

#Function will create a list of data frames, based on bootstrapping the initial data frame
#Iterations for the bootstrapping can be defined (default = 50)
bootstrapping_df_list_creator = function(df,
                                         iter = 50) {
  #the data frame is split into two separate DFs with NAs and without NAs
  df_impute_w_NA = df[!complete.cases(df),]
  df_impute_wo_NA = df[complete.cases(df),]
  #a list of data frames is created
  df_list = foreach(i = 1:iter) %do% {
    #bootstrap the two different subsets
    sampling_wo_NA = df_impute_wo_NA[sample(nrow(df_impute_wo_NA), 
                                            nrow(df_impute_wo_NA), 
                                            replace = TRUE),]
    sampling_NA = df_impute_w_NA [sample(nrow(df_impute_w_NA ),
                                         nrow(df_impute_w_NA ),
                                         replace = TRUE),]
    #make one data frame of the two subsets with NAs and w/o NAs, and remove duplicated rows
    sampling = rbind(sampling_NA,
                     sampling_wo_NA) %>%
      distinct()
    return(sampling)
  }
}

#Function will create a list of data frames, based on subsampling randomly from the initial data frame
#Iterations for the subsampling can be defined (default = 50), and proportion of rows to be subsampled (default = .8)
subsampling_df_list_creator = function(df, 
                                       subsample = 0.8,
                                       iter = 50) {
  #the data frame is split into two separate DFs with NAs and without NAs
  df_impute_w_NA = df[!complete.cases(df),]
  df_impute_wo_NA = df[complete.cases(df),]
  #a list of data frames is created
  df_list = foreach(i = 1:iter) %do% {
    #bootstrap the two different subsets
    sampling_wo_NA = df_impute_wo_NA[sample(nrow(df_impute_wo_NA), 
                                            subsample * nrow(df_impute_wo_NA), 
                                            replace = FALSE),]
    sampling_NA = df_impute_w_NA [sample(nrow(df_impute_w_NA),
                                         subsample * nrow(df_impute_w_NA),
                                         replace = FALSE),]
    #make one data frame of the two subsets with NAs and w/o NAs
    sampling = rbind(sampling_NA,
                     sampling_wo_NA)
    return(sampling)
  }
  return(df_list)
}

#Function that takes a list of data frames from the previous functions, 
#and applies several learners for NA imputation on each data frame within the list.
#Imputated data frames are subsequently combined and the mean value for each variable
#is determined. Returned is a dataframe with the ID and all variables
impute_values = function(df_list,
                         learner = c("regr.cubist",
                                     "regr.rpart",
                                     "regr.gbm",
                                     "regr.xgboost")) {
  #if the learner is set to regr.xgboost the hyperparameters are set to nrounds = 10,
  #eta = 0.3, max_depth = 10, and subsample = 0.5
  if (learner == "regr.xgboost") {
    learner = makeLearner("regr.xgboost")
    par_list = list(
      nrounds = 10,
      eta = 0.3,
      max_depth = 10,
      subsample = 0.5
    )
    tuned_learner = setHyperPars(learner,
                                 par.vals = par_list)
    imputeMethod = imputeLearner(tuned_learner)
  } else {
    imputeMethod = imputeLearner(learner)
  }
  df_list_imputed = map(df_list,
                        ~ {
                          df = .
                          imp = impute(df[, -1],
                                       classes = list(numeric = imputeMethod))
                          df_imp = imp$data %>%
                            as.data.frame(stringsAsFactors = FALSE) %>%
                            mutate(ID = df[, 1])
                          df_imp = df_imp[, c(5, 1:4)]
                          colnames(df_imp) = c("ID",
                                               "V1",
                                               "V2",
                                               "V3",
                                               "V4")
                          return(df_imp)
                        }) 
  df_list_reduced = df_list_imputed %>%
    reduce(full_join,
           by = "ID")
  colname_pattern = c("V1", "V2", "V3", "V4")
  means = lapply(colname_pattern,
                 function(x) {
                   apply(df_list_reduced[, grep(x, names(df_list_reduced))],
                         1,
                         mean,
                         na.rm = TRUE)
                 })
  result = do.call(cbind,
                   means) %>%
    as.data.frame(stringsAsFactors = FALSE) %>%
    mutate(ID = df_list_reduced[, 1]) %>%
    arrange(ID)
  return(result)
}

#Function that takes the imputed data frame and the original data frame,
#and adds columns if a given variable has been imputed or not.
add_imputation_info = function(imputed_df,
                               df) {
  df = df %>% arrange(.[,1])
  imput_info = foreach(i = 2:5,
                       .combine = cbind) %do% {
                         V_i = df %$% .[,i]
                         imput_info_i = map_chr(V_i,
                                                ~{value = ifelse(is.na(.), "yes", "no")})
                         return(imput_info_i)
                       } %>%
    as.data.frame(stringsAsFactors = FALSE) %>%
    mutate(ID = df[,1])
  imputed_df_final = merge(imputed_df,
                           imput_info,
                           by = "ID")
  colnames(imputed_df_final)[6:9] = c("V1_imp", "V2_imp", "V3_imp", "V4_imp")
  return(imputed_df_final)
}

#Function that takes a dataframe with imputed values and labels, and creates density plots
#If a tiff path is given, the function will export a TIFF
plot_imputation_distributions_density = function(imputed_df,
                                                 path_tiff = NA) {
  
  col_names = c("V1", "V2", "V3", "V4")
  #create the ggplots
  plot_list = foreach(i = seq_along(col_names)) %do% {
    plot_i = ggplot(imputed_df,
                    aes_string(col_names[[i]],
                               fill = paste(col_names[[i]],
                                            "imp",
                                            sep = "_"))) +
      geom_density(alpha = .2,
                   size = .3) +
      scale_fill_manual("imputed?", values = c("yes" = "orange",
                                               "no" = "grey")) +
      theme_linedraw()
  }
  #Export a TIFF if a path is given
  if (!is.na(path_tiff)) {
    tiff_path_final = paste(path_tiff,
                            deparse(substitute(df)),
                            ".tif",
                            sep = "")
    tiff(
      tiff_path_final,
      width = 6,
      height = 4,
      unit = "in",
      res = 300
    )
    plot(grid.arrange(plot_list[[1]],
                      plot_list[[2]],
                      plot_list[[3]],
                      plot_list[[4]]))
    dev.off()
  }
  #show the grid arrange
  plot(grid.arrange(plot_list[[1]],
                    plot_list[[2]],
                    plot_list[[3]],
                    plot_list[[4]]))
}

#Function that takes a dataframe with imputed values and labels, and creates histograms
#If a tiff path is given, the function will export a TIFF
plot_imputation_distributions_hist = function(imputed_df,
                                              path_tiff = NA) {
  col_names = c("V1", "V2", "V3", "V4")
  #create the ggplots
  plot_list = foreach(i = seq_along(col_names)) %do% {
    plot_i = ggplot(imputed_df,
                    aes_string(col_names[[i]],
                               fill = paste(col_names[[i]],
                                            "imp",
                                            sep = "_"))) +
      geom_histogram(bins = 50) +
      scale_fill_manual("imputed?", values = c("yes" = "orange",
                                               "no" = "grey")) +
      theme_linedraw()
  }
  #Export a TIFF if a path is given
  if (!is.na(path_tiff)) {
    tiff_path_final = paste(path_tiff,
                            deparse(substitute(df)),
                            ".tif",
                            sep = "")
    tiff(
      tiff_path_final,
      width = 6,
      height = 4,
      unit = "in",
      res = 300
    )
    plot(grid.arrange(plot_list[[1]],
                      plot_list[[2]],
                      plot_list[[3]],
                      plot_list[[4]]))
    dev.off()
  }
  plot(grid.arrange(plot_list[[1]],
                    plot_list[[2]],
                    plot_list[[3]],
                    plot_list[[4]]))
}

get_imputation_stats = function(df_imp) {
  col_names = c("V1", "V2", "V3", "V4")
  #iterate through the variables and provide median and quantiles + distance metric for
  #the imputed and original values
  results_df = foreach(i = seq_along(col_names),
                       .combine = rbind) %do% {
                         col_name_i = col_names[[i]]
                         col_name2_i = paste(col_name_i,
                                             "imp",
                                             sep = "_")
                         values_original = df_imp %>%
                           filter(.[, col_name2_i] == "no") %>%
                           summarise(
                             median = median(.[, col_name_i]),
                             up_quant = quantile(.[, col_name_i],
                                                 probs = 0.75),
                             low_quant = quantile(.[, col_name_i],
                                                  probs = 0.25)
                           )
                         values_imputed = df_imp %>%
                           filter(.[, col_name2_i] == "yes") %>%
                           summarise(
                             median_imp = median(.[, col_name_i]),
                             up_quant_imp = quantile(.[, col_name_i],
                                                     probs = 0.75),
                             low_quant_imp = quantile(.[, col_name_i],
                                                      probs = 0.25)
                           )
                         ks_difference = suppressWarnings(ks.test(
                           df_imp %>%
                             filter(.[, col_name2_i] == "yes") %$%
                             .[, col_name_i],
                           df_imp %>%
                             filter(.[, col_name2_i] == "no") %$%
                             .[, col_name_i]
                         ))
                         distance = as.numeric(ks_difference$statistic)
                         result = cbind(col_name_i,
                                        values_original,
                                        values_imputed,
                                        distance)
                       } %>%
    as.data.frame(stringsAsFactors = FALSE)
  colnames(results_df)[[1]] = "variable"
  return(results_df)
}

# Data --------------------------------------------------------------------

#Loads in the provided example data frame
df = read.csv(
  "~/Downloads/example_data.csv", #set the path of the example CSV
  stringsAsFactors = FALSE
)

#Create different lists of data frame based on a CV-type approach, bootstrapping and subsampling
df_list_CV = CV_df_list_creater(df)
df_list_BT = bootstrapping_df_list_creator(df)
df_list_Sub = subsampling_df_list_creator(df)

#Define the to be tested learner for imputation
#Example learners used in the publication
#"regr.cubist"
#"regr.rpart"
#"regr.gbm"
#"regr.xgboost"
to_test_learner = "regr.xgboost"

#Apply the learner on the different list of data frames
impute_CV = impute_values(df_list_CV,
                          to_test_learner)
impute_BT = impute_values(df_list_BT,
                          to_test_learner)
impute_Sub = impute_values(df_list_Sub,
                           to_test_learner)

#Add additional information if a given variable was imputed or not
impute_CV = add_imputation_info(impute_CV,
                                df)
impute_BT = add_imputation_info(impute_BT,
                                df)
impute_Sub = add_imputation_info(impute_Sub,
                                 df)

#Generate plots and a data frame with the statistics for the CV-type approach for multiple imputation
plot_imputation_distributions_density(impute_CV)
plot_imputation_distributions_hist(impute_CV)
statistics_CV = get_imputation_stats(impute_CV)

#Generate plots and a data frame with the statistics for the Bootstrapping approach for multiple imputation
plot_imputation_distributions_density(impute_BT)
plot_imputation_distributions_hist(impute_BT)
statistics_BT = get_imputation_stats(impute_BT)

#Generate plots and a data frame with the statistics for the Subsampling approach for multiple imputation
plot_imputation_distributions_density(impute_Sub)
plot_imputation_distributions_hist(impute_Sub)
statistics_Sub = get_imputation_stats(impute_Sub)
```
