## Supplementary Information for "*N*-myristoyltransferase inhibition is synthetic lethal in MYC-deregulated cancers"

**Affiliations**:

### Table of contents **Page**

| Cell line | IMP1031 | | IMP1036 | | IMP1088 | |
|---|---|---|---|---|---|---|
| | $EC_{50}$ [nM] | Max. effect [%] | $EC_{50}$ [nM] | Max. effect [%] | $EC_{50}$ [nM] | Max. effect [%] |
| BL41 | 38 | 97 | 155 | 99 | 21 | 100 |
| Ramos | 52 | 98 | 205 | 97 | 117 | 96 |
| Raji | 67 | 76 | 225 | 74 | 63 | 63 |
| CA46 | 70 | 66 | 249 | 75 | 43 | 72 |
| Farage | 180 | 70 | 552 | 63 | 38 | 98 |
| Jurkat | 102 | 97 | 478 | 95 | 19 | 95 |
| Karpas 422 | 44 | 90 | 171 | 89 | 13 | 82 |
| SU DHL8 | 222 | 33 | 323 | 34 | 84 | 43 |
| WSU NHL | 64 | 90 | 240 | 87 | 397 | 64 |
| HeLa | 55 | 93 | 292 | 78 | 147 | 100 |
| MDA-MB-231 | 55 | 90 | 98 | 87 | 21 | 94 |
| BL70 | *nd* | *nd* | *nd* | *nd* | 10 | 100 |
| Daudi | *nd* | *nd* | *nd* | *nd* | 52 | 76 |
| Namalwa | *nd* | *nd* | *nd* | *nd* | 22 | 47 |
| Rael | *nd* | *nd* | *nd* | *nd* | 40 | 88 |
| Riva | *nd* | *nd* | *nd* | *nd* | 73 | 80 |

**Supplementary Figure 1: Potency of NMT inhibitors IMP1031, IMP1036 and IMP1088 against various cancer cell lines.**

$EC_{50}$ for each inhibitor and the maximum effect (100% effect determined for 10 µg/mL puromycin control) after 72 hours, with the respective NMT inhibitor. Values for Jurkat, MDA-MB-231 and HeLa cells were calculated using data from *(1)*. nd: not determined.

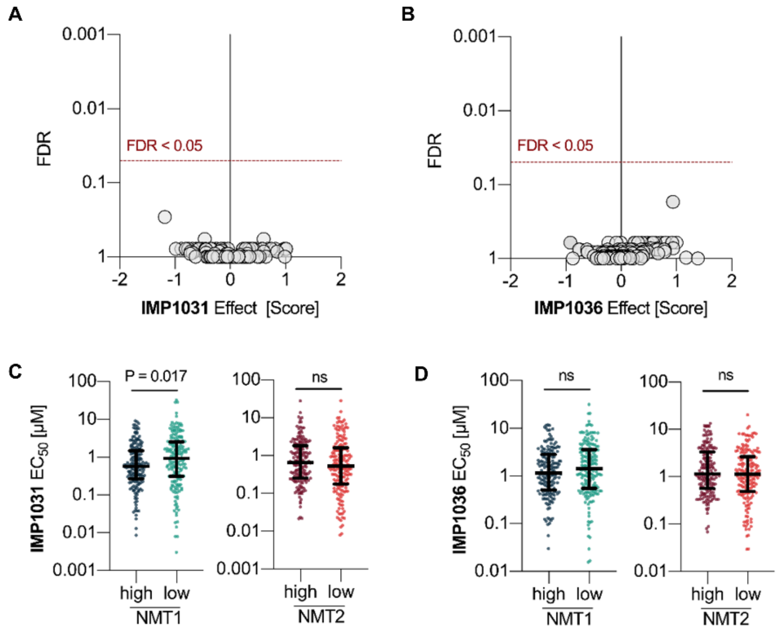

E

| Compound | Cancer functional event (CFE) | ∆Mean $EC_{50}$ | Adjusted p-value |
|---|---|---|---|
| **IMP1031** | Loss.cnaPANCAN381..ARFGAP3 | **0.8** | 0.11 |
| | MAP3K1_mut | **-1.49** | 0.12 |
| **IMP1036** | BBRM1_mut | **-0.58** | 0.23 |
| **IMP366 (ICL1100013)** | NA | NA | NA |

**Supplementary Figure 2: Mutation status of NMT substrates and expression levels of NMT1/2 are not predictive of sensitivity to NMT inhibition.**

**(A)** Volcano plot showing the effect of the presence of mutations in a given NMT substrate (n = 125) on sensitivity/resistance to IMP1031 and **(B)** IMP1036 (ANOVA test). **(C)** Cell lines screened against IMP1031 or **(D)** IMP1036 were divided by quantiles into high and low expressers of NMT1 or NMT2 and $EC_{50}$ values compared (ANOVA test). **(E)** Effect of CFEs on responsiveness to NMT inhibitors IMP1031, IMP1036 and IMP366 (published under the name ICL1100013 – extracted from *(2)*).

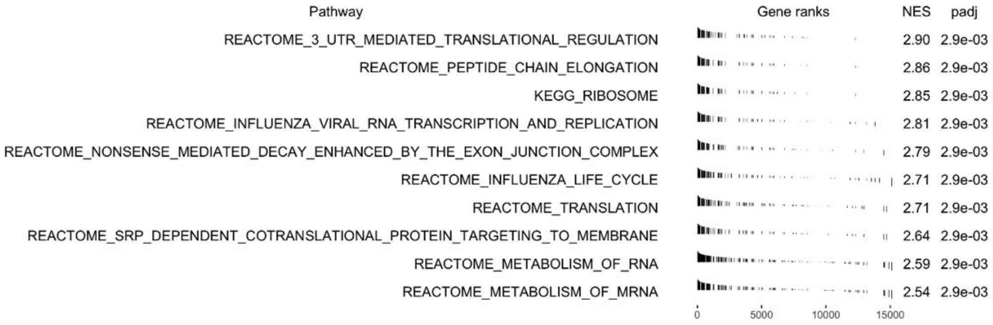

**Supplementary Figure 3: Gene sets enriched in cell lines sensitive to IMP1031.**

The top 10 upregulated annotated gene sets by normalized enrichment score (NES) in cancer cell lines sensitive to IMP1031 are shown.

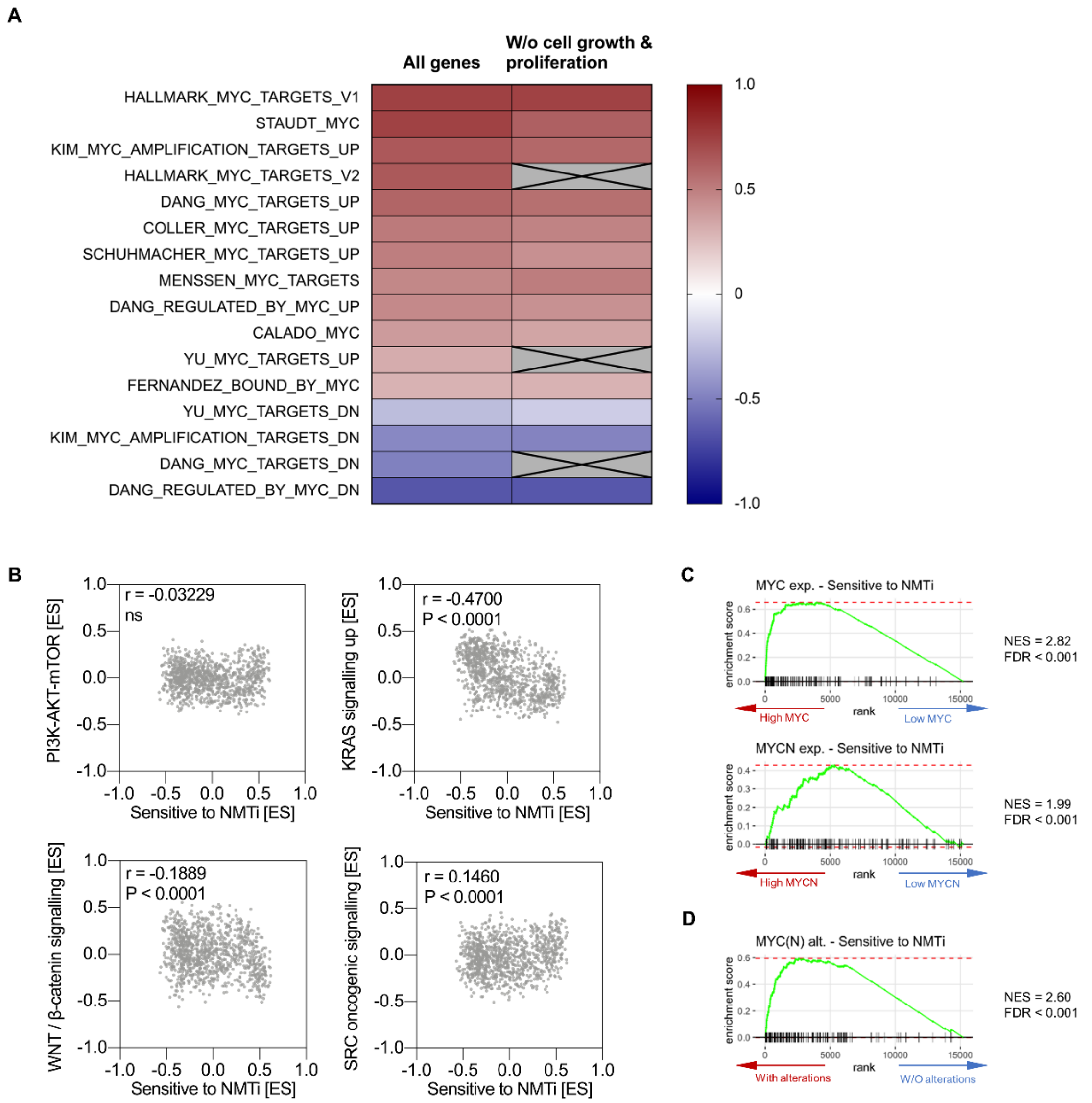

**Supplementary Figure 4: Enrichment of the ‘sensitive to NMTi’ gene set correlates with MYC dysregulation and expression, but not with other oncogenic signatures.**

**(A)** Spearman correlation coefficients of the ‘Sensitive to NMTi’ gene set with multiple MYC-related gene sets, performed either with all genes in the signature or after the removal of genes with GO terms ‘proliferation’, ‘growth’ and ‘cell cycle’ (greyed out variables indicate too few genes for reliable GSVA). **(B)** GSVA was performed on the COSMIC cell lines with gene sets related to oncogenic signaling and correlation of the ES for the ‘Sensitive to NMTi’ gene set was assessed (Spearman rank test). **(C)** Transcription in cell lines with increased MYC or MYCN expression is strongly enriched for the ‘Sensitive to NMTi’ gene set. **(D)** Transcription in cell lines with structural alterations in the MYC and/or MYCN loci is strongly enriched for the ‘Sensitive to NMTi’ gene set.

**A**

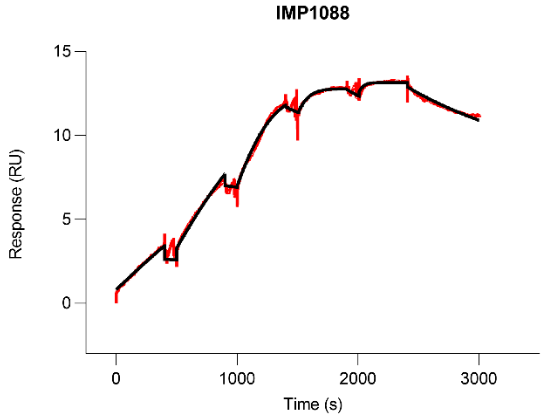

**B**

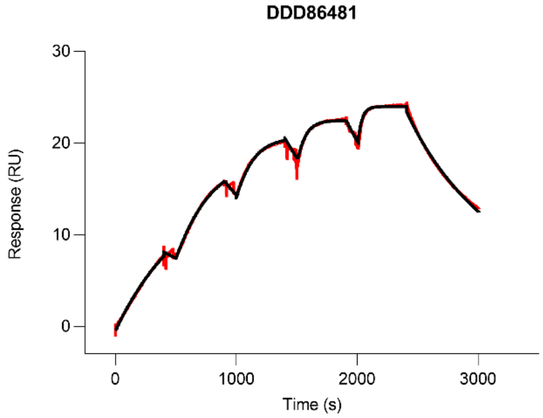

**C**

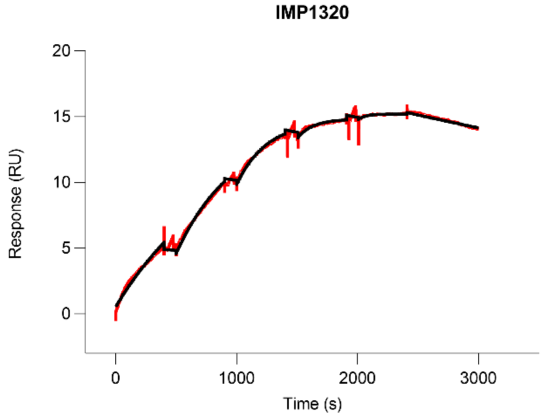

**D**

| Compound | $k_a$ (1/Ms) x10$^6$ | SEM | $k_d$ (1/s) x10$^{-3}$ | SEM | $K_D$ (nM) | SEM |
|---|---|---|---|---|---|---|
| IMP1088 | 3.37 | 0.10 | 0.35 | 0.01 | 0.10 | 0.01 |
| IMP1320 | 1.42 | 0.08 | 0.13 | 0.01 | 0.09 | 0.01 |
| DDD86481 | 2.93 | 0.08 | 1.28 | 0.06 | 0.43 | 0.01 |

**Supplementary Figure 5: Representative SPR sensorgrams of single cycle kinetics measurements of NMTi interacting with recombinant NMT1.**

**(A)** IMP1088. **(B)** DDD86481. **(C)** IMP1320. Sensorgrams were fitted to a 1:1 model, (black lines). **(D)** Kinetic constants for binding of compounds IMP1088, IMP1320 and DDD86481 to recombinant NMT1 protein using single cycle kinetics measurements. SEM - Standard error of the mean.

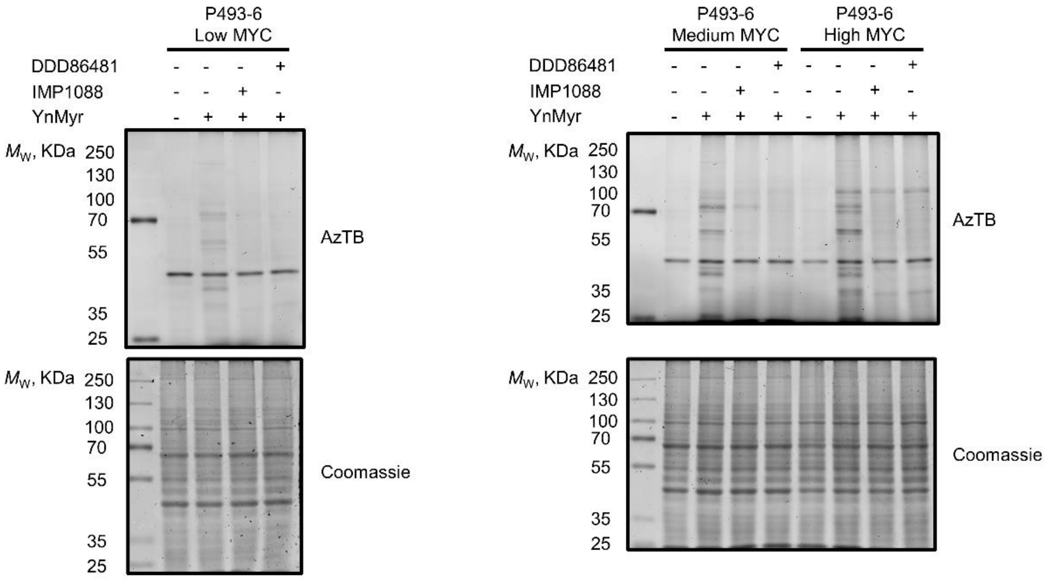

**Supplementary Figure 6: NMTi target engagement in P493-6 cells.**

In-gel fluorescence assay was performed on lysates from P493-6 cells (low, medium, high MYC) treated with IMP1088 (100 nM) or DDD86481 (1 µM) and YnMyr for 18 h and subjected to ligation with Azido-TAMRA-biotin (AzTB).

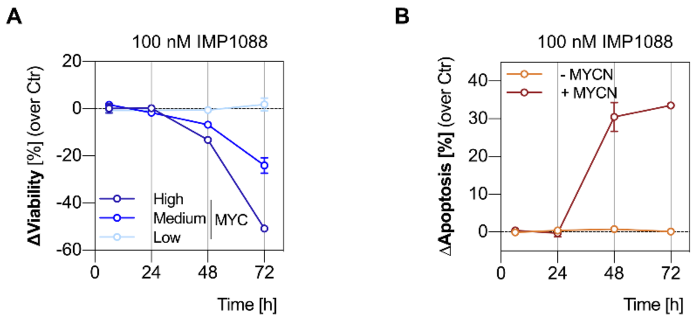

**Supplementary Figure 7: Impact of IMP1088 on viability and apoptosis in P493-6 and SHEP cell lines.**

**(A)** Cell viability of P493-6 cells with varied MYC levels in the presence of 100 nM IMP1088 compared to DMSO control, measured via membrane integrity over time using FACS. Data are shown as mean ± s.e.m. of n = 2 biological replicates. **(B)** Cell viability of SHEP cells with or without MYCN induction in the presence of 100 nM IMP1088 compared to DMSO control, measured via membrane integrity over time using FACS. Data are shown as mean ± s.e.m. of n = 2 biological replicates.

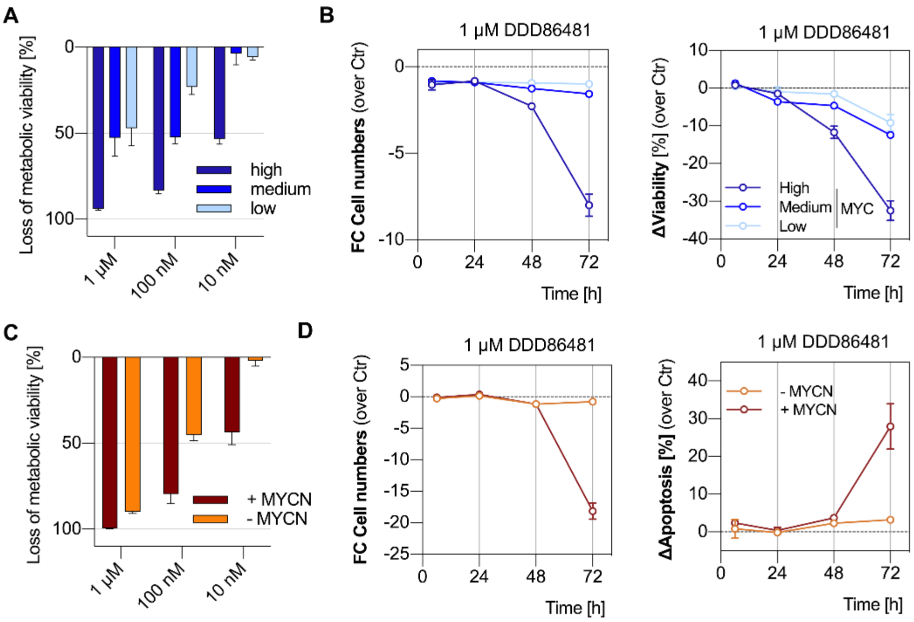

**Supplementary Figure 8: DDD86841 reproduces the synthetic lethality of NMTi upon MYC or MYCN induction.**

**(A)** Metabolic activity in low, medium and high MYC expressing P493-6 cell line upon NMT inhibition with DDD86841. Data are shown as mean ± s.e.m. of n = 4 biological replicates. **(B)** Fold-change in cell number, and cell viability measured via membrane integrity over time using FACS for P493-6 cell line with varied MYC levels comparing 1 µM DDD86841 versus DMSO control over time using FACS. Data are shown as mean ± s.e.m. of n = 2 biological replicates. **(C)** Metabolic activity in SHEP cell line with or without MYCN induction upon NMT inhibition with DDD86841. Data are shown as mean ± s.e.m. of n = 4 biological replicates. **(D)** Fold-change in cell number and decrease in cell viability measured via membrane integrity over time using FACS for SHEP cell line with or without MYCN induction comparing 1 µM DDD86841 versus DMSO control. Data are shown as mean ± s.e.m. of n = 2 biological replicates.

| Tumor model | | rel. $IC_{50}$ (µM) | t | b | abs. $IC_{50}$ (µM) | t | b | rel. $IC_{70}$ (µM) | t | b | abs. $IC_{70}$ (µM) | t | b |
|---|---|---|---|---|---|---|---|---|---|---|---|---|---|
| BXF | 1258 | 0.680 | 95 | 1 | 0.655 | 95 | 1 | 0.995 | 95 | 1 | 0.981 | 95 | 1 |
| BXF | 2775 | 2.847 | 87 | 0 | 2.544 | 87 | 0 | 3.940 | 87 | 0 | 3.643 | 87 | 0 |
| CEXF | 773 | 1.575 | 106 | 0 | 1.669 | 106 | 0 | 2.510 | 106 | 0 | 2.620 | 106 | 0 |
| CNXF | 2601 | 0.441 | 97 | 0 | 0.429 | 97 | 0 | 0.708 | 97 | 0 | 0.693 | 97 | 0 |
| CNXF | 2615 | 0.186 | 98 | 32 | 0.666 | 98 | 32 | 0.575 | 98 | 32 | 10.000 | 98 | 32 |
| CXA | 3113 | 1.548 | 110 | 8 | 1.806 | 110 | 8 | 2.261 | 110 | 8 | 2.750 | 110 | 8 |
| CXF | 1086 | 0.982 | 92 | 5 | 0.968 | 92 | 5 | 1.208 | 92 | 5 | 1.230 | 92 | 5 |
| CXF | 2083 | 1.850 | 102 | 2 | 1.926 | 102 | 2 | 2.829 | 102 | 2 | 2.973 | 102 | 2 |
| CXF | 280 | 0.576 | 92 | 0 | 0.434 | 92 | 0 | 2.300 | 92 | 0 | 1.861 | 92 | 0 |
| CXF | 742 | 1.038 | 95 | 25 | 1.507 | 95 | 25 | 1.755 | 95 | 25 | 5.451 | 95 | 25 |
| CXF | RKO (L) | 0.066 | 98 | 0 | 0.065 | 98 | 0 | 0.096 | 98 | 0 | 0.095 | 98 | 0 |
| CXF | RKO | 0.101 | 98 | 13 | 0.113 | 98 | 13 | 0.152 | 98 | 13 | 0.197 | 98 | 13 |
| HNXF | 1838 | 0.597 | 90 | 0 | 0.530 | 90 | 0 | 0.953 | 90 | 0 | 0.876 | 90 | 0 |
| HNXF | 908 | 0.424 | 93 | 0 | 0.354 | 93 | 0 | 1.230 | 93 | 0 | 1.077 | 93 | 0 |
| LIXFH | 658 | 0.213 | 90 | 0 | 0.178 | 90 | 0 | 0.434 | 90 | 0 | 0.381 | 90 | 0 |
| LXFL | 1121 | 0.093 | 108 | 2 | 0.110 | 108 | 2 | 0.206 | 108 | 2 | 0.243 | 108 | 2 |
| LXFL | 1674 | 0.112 | 102 | 2 | 0.116 | 102 | 2 | 0.177 | 102 | 2 | 0.187 | 102 | 2 |
| LXFL | 529 (L) | 1.169 | 96 | 6 | 1.259 | 96 | 6 | 3.715 | 96 | 6 | 4.807 | 96 | 6 |
| LXFL | 529 | 0.595 | 99 | 3 | 0.611 | 99 | 3 | 1.033 | 99 | 3 | 1.099 | 99 | 3 |
| LXFS | 650 | 0.022 | 105 | 0 | 0.022 | 105 | 0 | 0.027 | 105 | 0 | 0.027 | 105 | 0 |
| LYXFDLBC | 2835 | 0.055 | 90 | 0 | 0.047 | 90 | 0 | 0.108 | 90 | 0 | 0.096 | 90 | 0 |
| MAXFTN | 401 | 1.943 | 104 | 0 | 2.110 | 104 | 0 | 5.408 | 104 | 0 | 5.756 | 104 | 0 |
| MAXFTN | 449 | 0.565 | 97 | 0 | 0.543 | 97 | 0 | 1.004 | 97 | 0 | 0.976 | 97 | 0 |
| MAXFTN | 574 | 0.246 | 100 | 1 | 0.252 | 100 | 1 | 0.581 | 100 | 1 | 0.603 | 100 | 1 |
| MAXFTN | 583 | 1.833 | 87 | 0 | 1.555 | 87 | 0 | 2.958 | 87 | 0 | 2.637 | 87 | 0 |
| MAXFTN | 857 | 0.403 | 97 | 0 | 0.389 | 97 | 0 | 0.619 | 97 | 0 | 0.603 | 97 | 0 |
| MEXF | 1792 | 0.083 | 95 | 28 | 0.209 | 95 | 28 | 0.251 | 95 | 28 | 10.000 | 95 | 28 |
| MEXF | 535 | 0.114 | 107 | 3 | 0.133 | 107 | 3 | 0.226 | 107 | 3 | 0.269 | 107 | 3 |
| MEXF | 566 | 2.497 | 93 | 32 | 5.318 | 93 | 32 | 5.124 | 93 | 32 | 10.000 | 93 | 32 |
| MEXF | 622 | 0.114 | 104 | 1 | 0.138 | 104 | 1 | 0.718 | 104 | 1 | 0.862 | 104 | 1 |
| OEXF | 2417 | 0.230 | 86 | 1 | 0.191 | 86 | 1 | 0.391 | 86 | 1 | 0.349 | 86 | 1 |
| OVXF | 1023 | 0.138 | 95 | 0 | 0.126 | 95 | 0 | 0.300 | 95 | 0 | 0.280 | 95 | 0 |
| OVXF | 1353 | 3.052 | 94 | 14 | 3.216 | 94 | 14 | 3.801 | 94 | 14 | 4.373 | 94 | 14 |
| OVXF | 1544 | 8.651 | 96 | 30 | 10.000 | 96 | 30 | 10.000 | 96 | 30 | 10.000 | 96 | 30 |
| OVXF | 1993 | 2.084 | 100 | 0 | 2.065 | 100 | 0 | 6.315 | 100 | 0 | 6.273 | 100 | 0 |
| OVXF | 899 | 1.794 | 96 | 0 | 1.656 | 96 | 0 | 4.219 | 96 | 0 | 3.978 | 96 | 0 |
| OVXF | OV-028 | 1.059 | 112 | 4 | 1.088 | 112 | 4 | 1.146 | 112 | 4 | 1.176 | 112 | 4 |
| PAXF | 1881 | 10.000 | | | 10.000 | | | 10.000 | | | 10.000 | | |
| PAXF | 1900 | 0.709 | 107 | 8 | 0.845 | 107 | 8 | 1.154 | 107 | 8 | 1.463 | 107 | 8 |
| PAXF | 1965 | 4.081 | 92 | 2 | 3.744 | 92 | 2 | 6.843 | 92 | 2 | 6.583 | 92 | 2 |
| PAXF | 1986 | 1.540 | 99 | 0 | 1.527 | 99 | 0 | 2.125 | 99 | 0 | 2.118 | 99 | 0 |
| PAXF | 1997 | 1.307 | 102 | 7 | 1.478 | 102 | 7 | 2.281 | 102 | 7 | 2.759 | 102 | 7 |
| PRXF | MRI-H-1579 | 2.584 | 95 | 0 | 2.425 | 95 | 0 | 4.166 | 95 | 0 | 3.980 | 95 | 0 |
| RXA | SMTCA75 | 6.144 | 94 | 0 | 4.792 | 94 | 0 | 10.000 | 94 | 0 | 10.000 | 94 | 0 |
| RXF | 2502 | 2.767 | 98 | 13 | 3.173 | 98 | 13 | 4.312 | 98 | 13 | 5.746 | 98 | 13 |
| RXF | 2667 | 10.000 | 101 | 20 | 10.000 | 101 | 20 | 10.000 | 101 | 20 | 10.000 | 101 | 20 |
| RXF | 2717 | 2.094 | 100 | 15 | 2.367 | 100 | 15 | 2.755 | 100 | 15 | 3.502 | 100 | 15 |
| RXF | 486 | 4.425 | 104 | 30 | 4.922 | 104 | 30 | 4.857 | 104 | 30 | 7.736 | 104 | 30 |
| SXA | SMTCA96 | 0.365 | 95 | 0 | 0.343 | 95 | 0 | 0.585 | 95 | 0 | 0.560 | 95 | 0 |
| SXFO | 1410 | 0.138 | 99 | 1 | 0.140 | 99 | 1 | 0.252 | 99 | 1 | 0.260 | 99 | 1 |
| SXFS | HT-1080 (L) | 0.042 | 111 | 2 | 0.045 | 111 | 2 | 0.054 | 111 | 2 | 0.058 | 111 | 2 |
| UXF | 2890 | 0.030 | 100 | 1 | 0.031 | 100 | 1 | 0.083 | 100 | 1 | 0.086 | 100 | 1 |
| **Mean $IC_{50}$ (geometr.)** | | 0.630 | | | 0.670 | | | 1.121 | | | 1.365 | | |

**Supplementary Figure 9: Potency of the NMT inhibitor IMP1320 measured in clonogenic analysis of 3D-cultured Patient-Derived cells.**

The relative $IC_{50}$ is determined as the concentration that gives a response halfway between the maximal signal (top plateau) and the maximally inhibited signal (bottom plateau) and equal to the inflection point of the sigmoidal concentration effect curve. The absolute $IC_{50}$ is determined as the concentration at the intersection of the concentration effect curve with T/C = 50%. If T/C<50% (T/C>75%) was detected for all test concentrations, the lowest (highest) concentration is given and used for calculation of the mean $IC_{50}$ value. t,b: top, bottom plateau of concentration-effect curve n.d.: not determined.

A

**Low expression**

| Cluster | Term | Count | FDR |
|---|---|---|---|
| 1 | Integral component of membrane | 70 | 0.6 |
| 2 | cadherin binding involved in cell-cell adhesion | 10 | 2 |
| 5 | cell adhesion | 12 | 5 |

**High expression**

| Cluster | Term | Count | FDR |
|---|---|---|---|
| 1 | RNA splicing | 11 | 0.1 |
| 2 | cell division | 16 | 0.1 |
| 3 | mRNA splicing, via spliceosome | 18 | 9.0477E-06 |
| 4 | spindle pole | 11 | 0.00177695 |
| 5 | mRNA export from nucleus | 10 | 0.01 |
| 8 | DNA repair | 11 | 1.7 |
| 11 | DNA binding | 46 | 0.00431931 |
| 12 | catalytic step 2 spliceosome | 6 | 8.2 |
| 13 | protein export from nucleus | 5 | 0.94 |

B

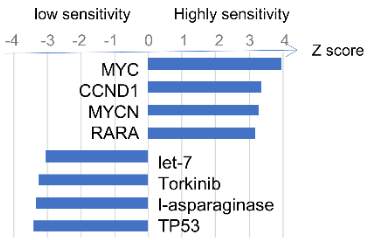

C

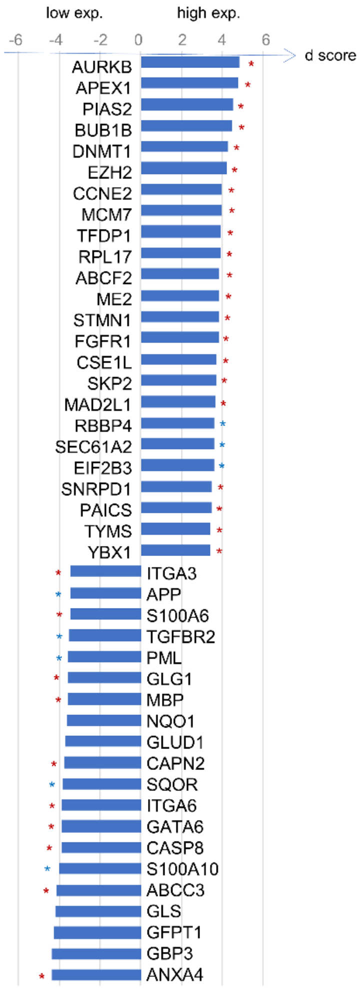

**Supplementary Figure 10: Functional analysis performed on positively and negatively correlated genes to the $log_{10}IC_{50}$.**

**(A)** Results of the analysis with the web-based tool DAVID for genes with a lower (top table) or higher (bottom table) expression in highly sensitive PD cancer cells. The number of genes present in the term is reported in count. FDR: false discovery rate. **(B)** Z score for each significant factor in the IPA analysis. A positive Z score indicates a factor that is predictive to be active in highly sensitive PD cancer cell, whereas a negative Z score indicate a factor that is active in PD cancer cells showing poor sensitivity to NMT inhibition. MYC was found as the highest scoring signature in highly sensitive tumors. **(C)** Targets of MYC that are correlated to $IC_{50}$ represented in the IPA knowledge base. The plot reports the d statistics from the quantitative SAM analysis (d score) as well as whether the direction of observed correlation is consistent with the expected direction in the knowledge base (red asterisk) or whether it is recognized as a MYC target, but the direction is not known in the database (blue asterisk).

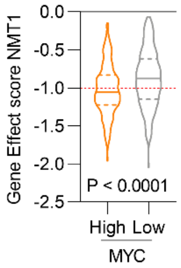

**Supplementary Figure 11: High dependence on NMT1 is correlated with elevated MYC expression.**

Cancer cell lines with highest MYC expression have lower gene effect scores of NMT1 i.e., greater dependence on NMT1 for optimal proliferation/viability compared to cell lines with lowest MYC expression (by quantiles; Wilcoxon rank test).

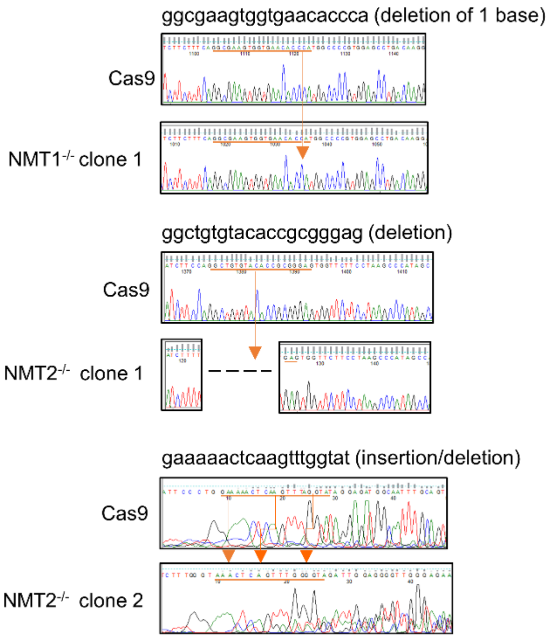

**Supplementary Figure 12: Representative chromatograms showing indels in CRISPR-Cas9 mediated NMT1 and NMT2 knockout in HeLa cells.**

Representative chromatograms of PCR products from wild type and mutant alleles, including insertions and/or deletion, in the modified HeLa cell line. The sequence of the gRNA primer is shown on top of the chromatogram and targeted regions are highlighted by arrows.

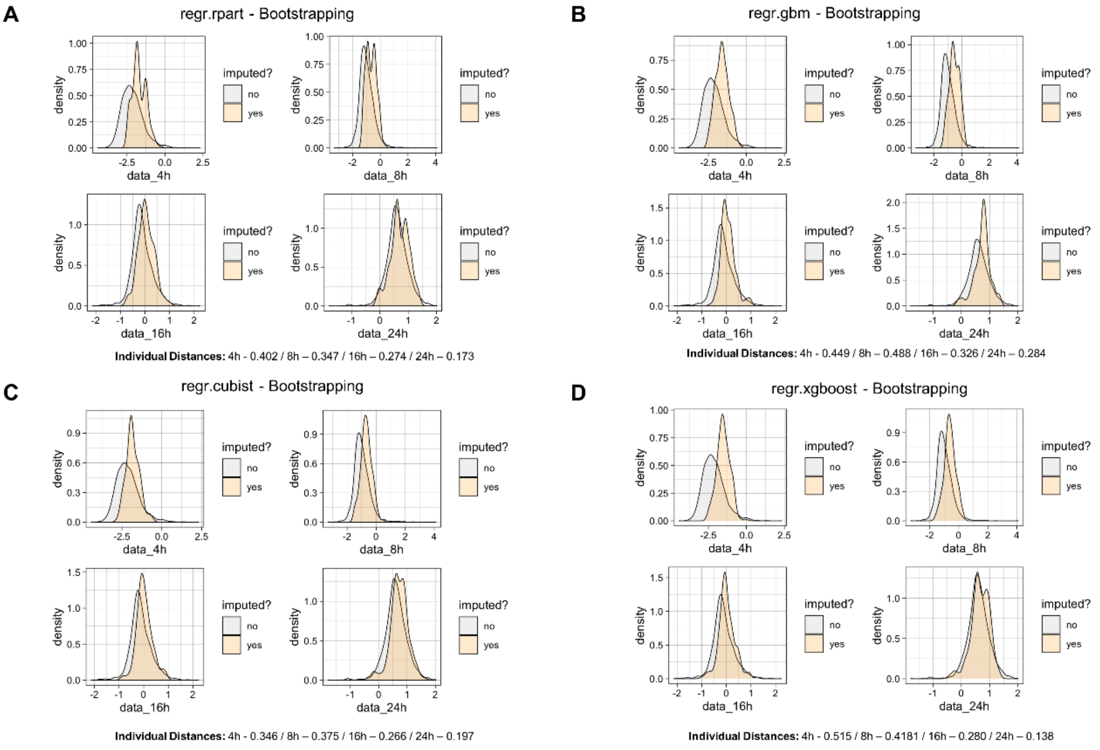

**E**

| Learner | CV-like approach | Bootstrapping | Resampling |
|---|---|---|---|
| regr.rpart | 1.427 | 1.205 | 1.387 |
| regr.xgboost | 1.241 | 1.362 | 1.304 |
| regr.gbm | 1.535 | 1.547 | 1.534 |
| regr.cubist | 1.186 | **1.184** | 1.199 |

**Supplementary Figure 13: Effect of different imputation learners and multiple imputation strategies**

**(A)** Example density plots, comparing the distribution of the imputed and none-imputed values for each time point, calculated for the regr.rpart learner with 50-fold bootstrapping on the data for multiple imputations. Density metrics per data point are based on Kolmogorov-Smirnov statistics, comparing the two distributions for each data point. **(B)** Example density plots, comparing the distribution of the imputed and none-imputed values for each time point, calculated for the regr.gbm learner with 50-fold bootstrapping on the data for multiple imputations. **(C)** Example density plots, comparing the distribution of the imputed and none-imputed values for each time point, calculated for the regr.cubist learner with 50-fold bootstrapping on the data for multiple imputations. **(D)** Example density plots, comparing the distribution of the imputed and none-imputed values for each time point, calculated for the regr.xgboost learner with 50-fold bootstrapping on the data for multiple imputations. **(E)** Table depicting the sum of the distance metrics across the four time points, based on Kolmogorov-Smirnov statistics, for every tested learner and multiple imputation strategy.

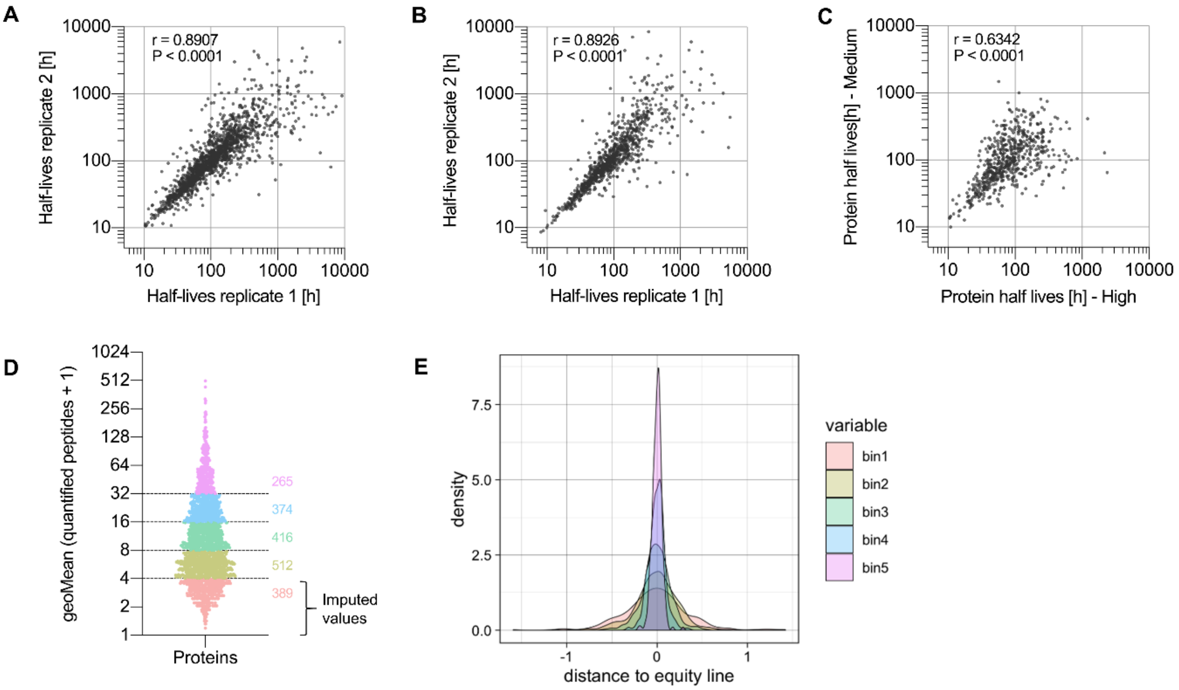

**Supplementary Figure 14: Reproducibility of calculated half-lives and effect of the binning strategy on proteome dynamics.**

**(A)** Scatterplot comparing replicate 1 and replicate 2 for the half-life calculations in the high-MYC condition. **(B)** Scatterplot comparing replicate 1 and replicate 2 for the half-life calculations in the medium-MYC condition. **(C)** Scatterplot showing the correlation of the average, overlapping half-lives as shown in Fig. 3B. **(D)** Example of binning strategy for high-MYC synthesis after 24 hours in the presence of 100 nM IMP1088 versus DMSO, on the geometric mean of the quantified spectra per data point. **(E)** Example density plot, for the same data point, showing the density per bin and distance to the equity line. (Spearman rank correlations – panels **A** to **C**.)

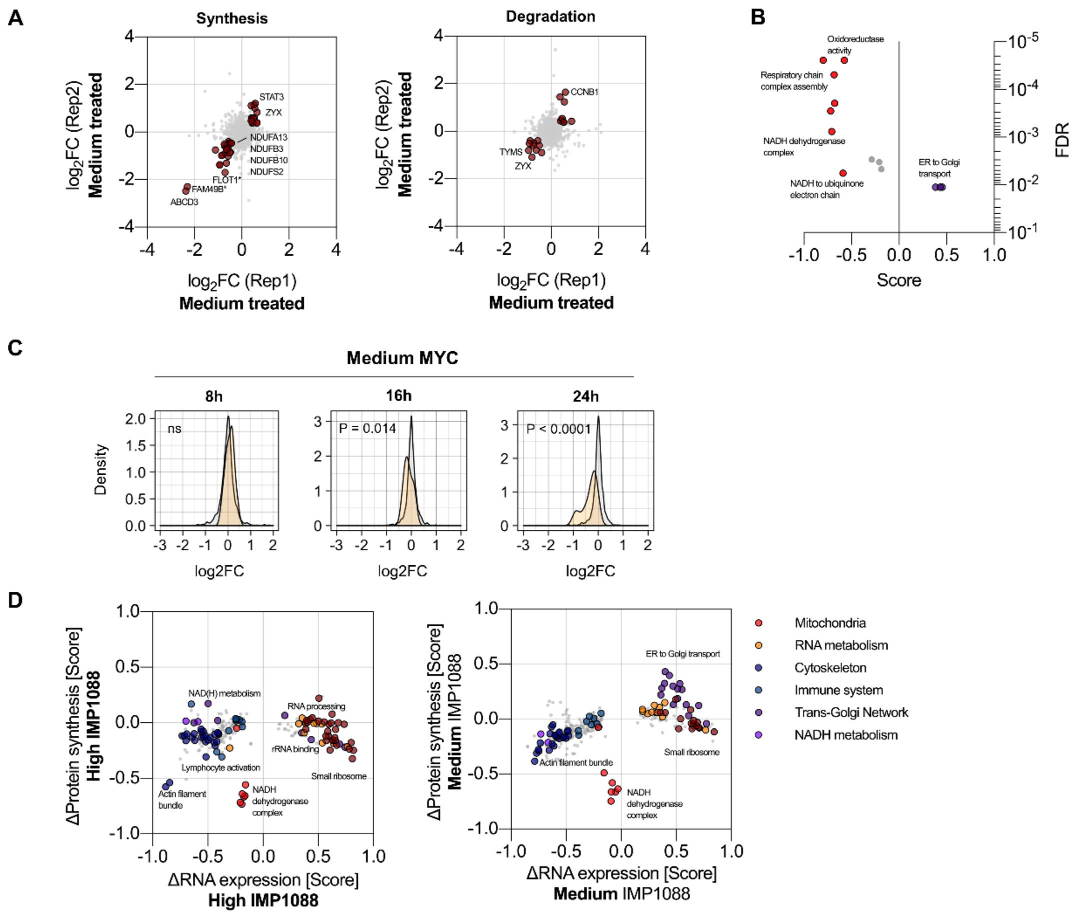

**Supplementary Figure 15: Impact of IMP1088 on protein synthesis and degradation.**

**(A)** Effect of IMP1088 on rates of protein synthesis (H/M SILAC ratio) and degradation of pre-existing proteins (M/L SILAC ratio) in medium-MYC cells (asterisk denotes NMT substrate). **(B)** 1D-enrichment on changes in synthesis rate for medium MYC; FDR threshold ≤ 2%. **(C)** Effect of IMP1088 on the synthesis of proteins of mitochondrial complex I over time in medium-MYC cells (Wilcoxon rank test). **(D)** 2D-enrichment between mRNA abundance changes and protein synthesis between control (DMSO) and 24 hours treatment with 100 nM IMP1088 for high and medium MYC; FDR threshold ≤ 0.1%.

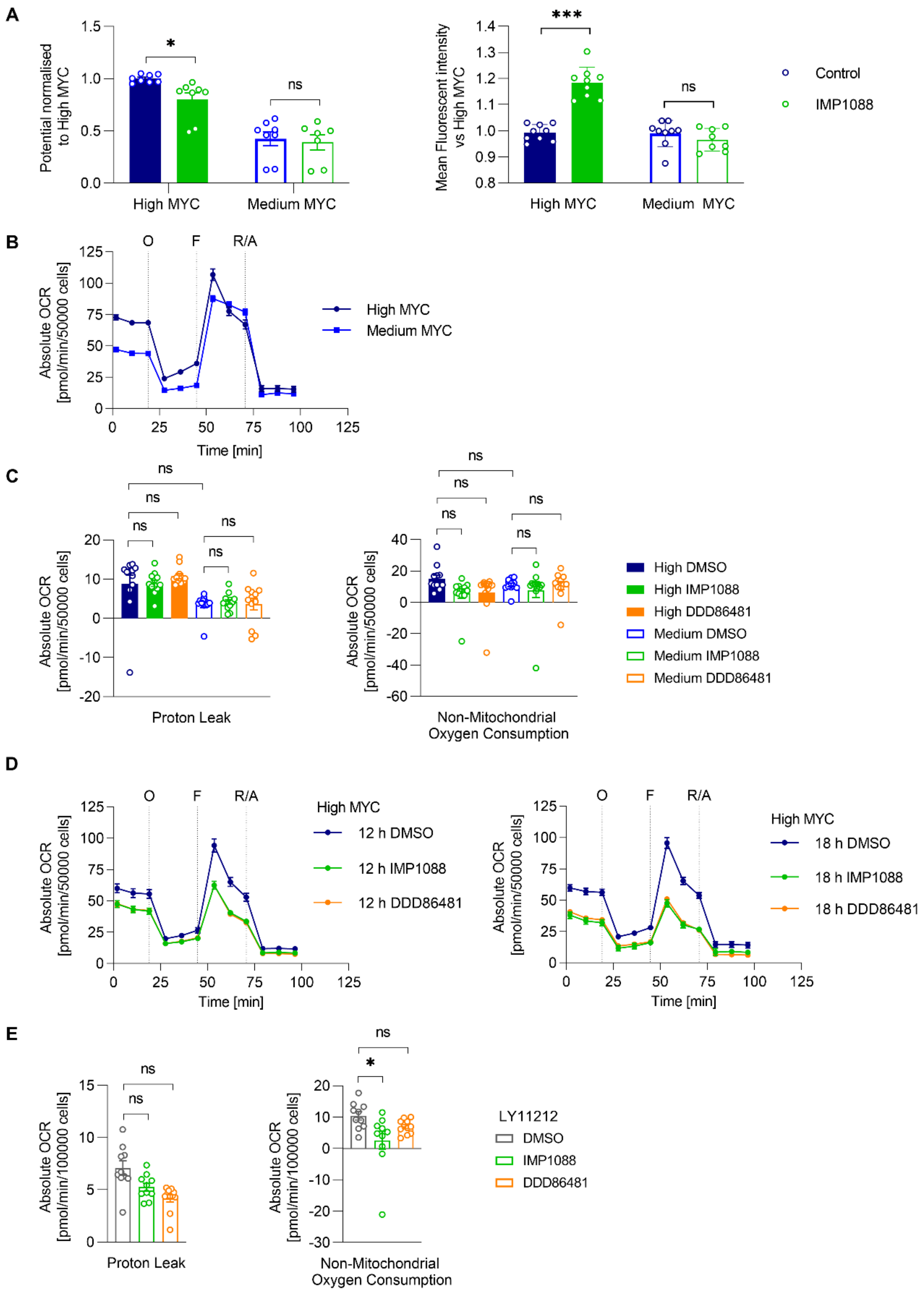

**Supplementary Figure 16: Impact of NMTi on mitochondrial function.**

**(A)** Left: Impact on mitochondrial potential by IMP1088 (100 nM, 18 h) and (right) superoxide production in high and medium MYC expressing P493-6 cells. Data are shown as mean ± s.e.m. of n = 3 biological replicates. **(B)** OCR of P493-6 cells expressing high or medium MYC, with DMSO vehicle treatment. **(C)** Parameters of mitochondrial function, proton leak and non-mitochondrial respiration, in P493-6 cells calculated using data from Fig. 4A.

**(D)** OCR of P493-6 cells expressing high MYC levels upon treatment with IMP1088 (100 nM) or DDD86481 (1 µM) for 12 or 18 h. **(E)** Parameters of mitochondrial function, proton leak and non-mitochondrial respiration, in LY11212 cells calculated using data from Fig. 4C. DMSO was used as a vehicle treatment. O: oligomycin, F: FCCP, R/A: rotenone and antimycin A. Data are shown as mean ± s.e.m. of $n = 3$ biological replicates ns: not statistically significant, * $P<0.05$ (two-way ANOVA).

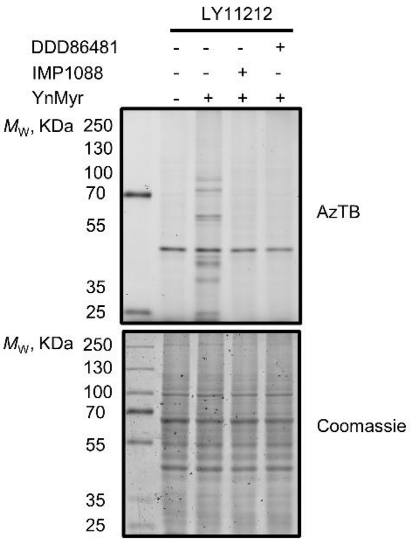

**Supplementary Figure 17: NMTi target engagement for LY11212.**

In-gel fluorescence assay performed on lysates from LY11212 cells treated with IMP1088 (100 nM) or DDD86481 (1 µM) and YnMyr for 18 h and subjected to ligation with Azido-TAMRA-biotin (AzTB).

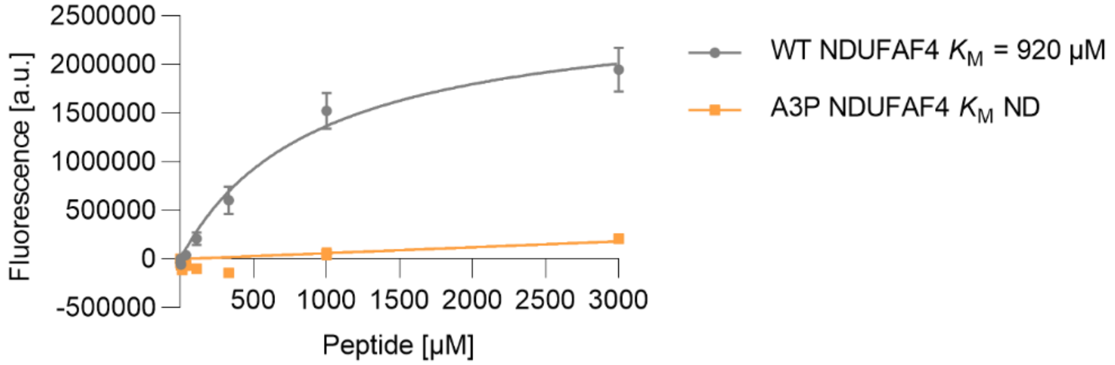

**Supplementary Figure 18: Calculation of $K_M$ for NMT activity in wild-type or A3P NDUFAF4 peptides.**

Synthetic NDUFAF4 (aa 2-10) peptides (WT and A3P) were subjected to NMT CPM assay. $K_M$ was determined using Michaelis Menten kinetics nonlinear regression fit with Prism (GraphPad). Data are shown as mean ± s.e.m. of n = 3 independent experiments. ND – not determined.

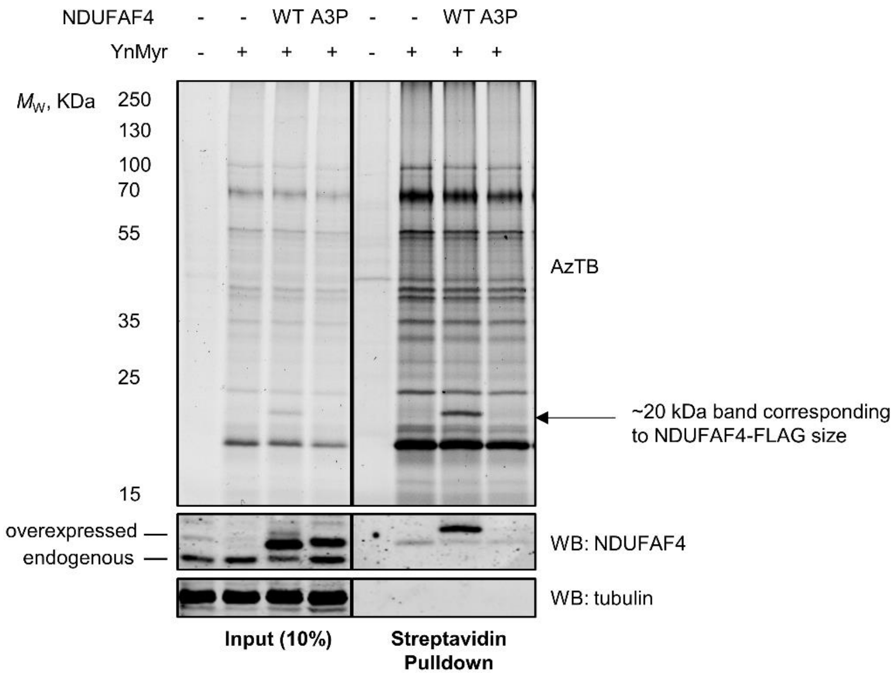

**Supplementary Figure 19: Validation of the lack of myristoylation of the NDUFAF4 A3P mutant.**

FLAG-tagged NDUFAF4 was overexpressed in HEK293 cells, which were treated with YnMyr 24 h post-transfection for a further 18 h. Cells were lysed, followed by CuAAC reaction and streptavidin pulldown.

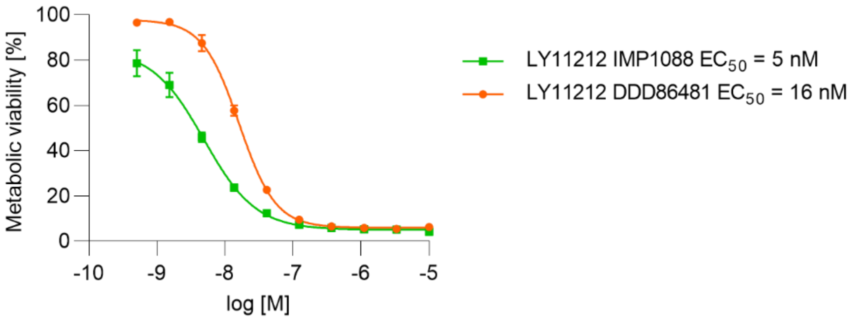

**Supplementary Figure 20: In-cell dose response of IMP1088 and DDD86481 in LY11212 cells.**

Metabolic viability was measured using the CellTiter-Blue assay on LY11212 PD cancer cells treated with a range of concentrations of IMP1088 and DDD86481 for 72 h. Data are shown as mean ± s.e.m. of n = 3 biological replicates.

| IP PK results | | | | | | | |
|---|---|---|---|---|---|---|---|
| Time (h) | Plasma levels (ng/ml) DDD86481 after IP dosing at 25 mg/kg | | | | | | |
| | M7 | M8 | M9 | Mean | ± | SD | CV (%) |
| **Body Weight (g)** | **31** | **32** | **32** | **32** | **±** | **0.58** | **1.82** |
| 0.00 | 0.0 | 0.0 | 0.0 | **0.0** | ± | **0.0** | **NA** |
| 0.08 | 23800.0 | 28900.0 | 36100.0 | **29600.0** | ± | **6179.8** | **20.9** |
| 0.25 | 56800.0 | 69200.0 | 61400.0 | **62466.7** | ± | **6268.4** | **10.0** |
| 0.50 | 59000.0 | 62400.0 | 54200.0 | **58533.3** | ± | **4119.9** | **7.0** |
| 1.00 | 53500.0 | 66600.0 | 51800.0 | **57300.0** | ± | **8098.8** | **14.1** |
| 2.00 | 40000.0 | 47800.0 | 37200.0 | **41666.7** | ± | **5493.0** | **13.2** |
| 4.00 | 39300.0 | 55100.0 | 42300.0 | **45566.7** | ± | **8391.3** | **18.4** |
| 8.00 | 13000.0 | 20400.0 | 13000.0 | **15466.7** | ± | **4272.4** | **27.6** |
| 24.00 | 24.6 | 18.8 | 14.0 | **19.1** | ± | **5.3** | **27.7** |

| IP PK Parameters | M7 | M8 | M9 | Mean | ± | SD | CV (%) |
|---|---|---|---|---|---|---|---|
| Nominal dose (mg/kg) | 25.0 | 25.0 | 25.0 | **25.0** | ± | **NA** | **NA** |
| No. points used for $t_{1/2}$ | 3 | 3 | 3 | **NA** | ± | **NA** | **NA** |
| Time points (hr) for $t_{1/2}$ | 4-24 | 4-24 | 4-24 | **NA** | ± | **NA** | **NA** |
| R-squared | 0.997 | 0.994 | 0.997 | **NA** | ± | **NA** | **NA** |
| Last time point (hr) for $AUC_{0\text{-}last}$ | 24.0 | 24.0 | 24.0 | **NA** | ± | **NA** | **NA** |
| Cmax (ng/ml) | 59000.0 | 69200.0 | 61400.0 | **63200.0** | ± | **5332.9** | **8.4** |
| Tmax (hr) | 0.5 | 0.25 | 0.25 | **0.3** | ± | **0.1** | **43.3** |
| AUC0-last (ng.hr/ml) | 304221.8 | 404005.4 | 303904.6 | **337377.3** | ± | **57701.8** | **17.1** |
| AUC0-inf (ng.hr/ml) | 304287.3 | 404051.2 | 303938.9 | **337425.8** | ± | **57699.5** | **17.1** |
| AUC0-inf/AUC0-last(%) | 100.02 | 100.01 | 100.01 | **100.0** | ± | **0.0** | **0.01** |
| Cl_F_obs (mL/min/kg) | 1.37 | 1.03 | 1.37 | **1.3** | ± | **0.2** | **15.6** |
| T1/2elim (hr) | 1.85 | 1.69 | 1.70 | **1.7** | ± | **0.1** | **5.0** |
| MRTlast (hr) | 4.01 | 4.22 | 3.97 | **4.1** | ± | **0.1** | **3.3** |
| MRTINF_obs (hr) | 4.01 | 4.22 | 3.97 | **4.1** | ± | **0.1** | **3.3** |
| %BA* | 158.33 | 210.24 | 158.15 | **175.6** | ± | **30.0** | **17.1** |

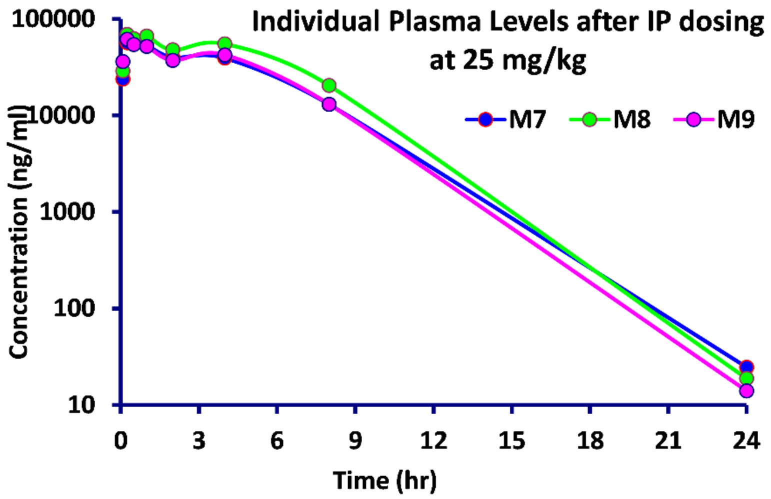

**Supplementary Figure 21: Pharmacokinetic profile of DDD86481.**

DDD86481 was dosed once at 25mg/kg interperitoneally (IP, N = 3, mice M7-M9), and plasma levels measured by high performance liquid chromatography-mass spectrometry (HPLC-MS) at the time points indicated. The concentration of DDD86481 remains above 1 µM (a concentration sufficient to fully suppress NMT activity in cells in culture) for ca. 16 hours post dosing, dropping to ca. 30 nM at 24 hours.

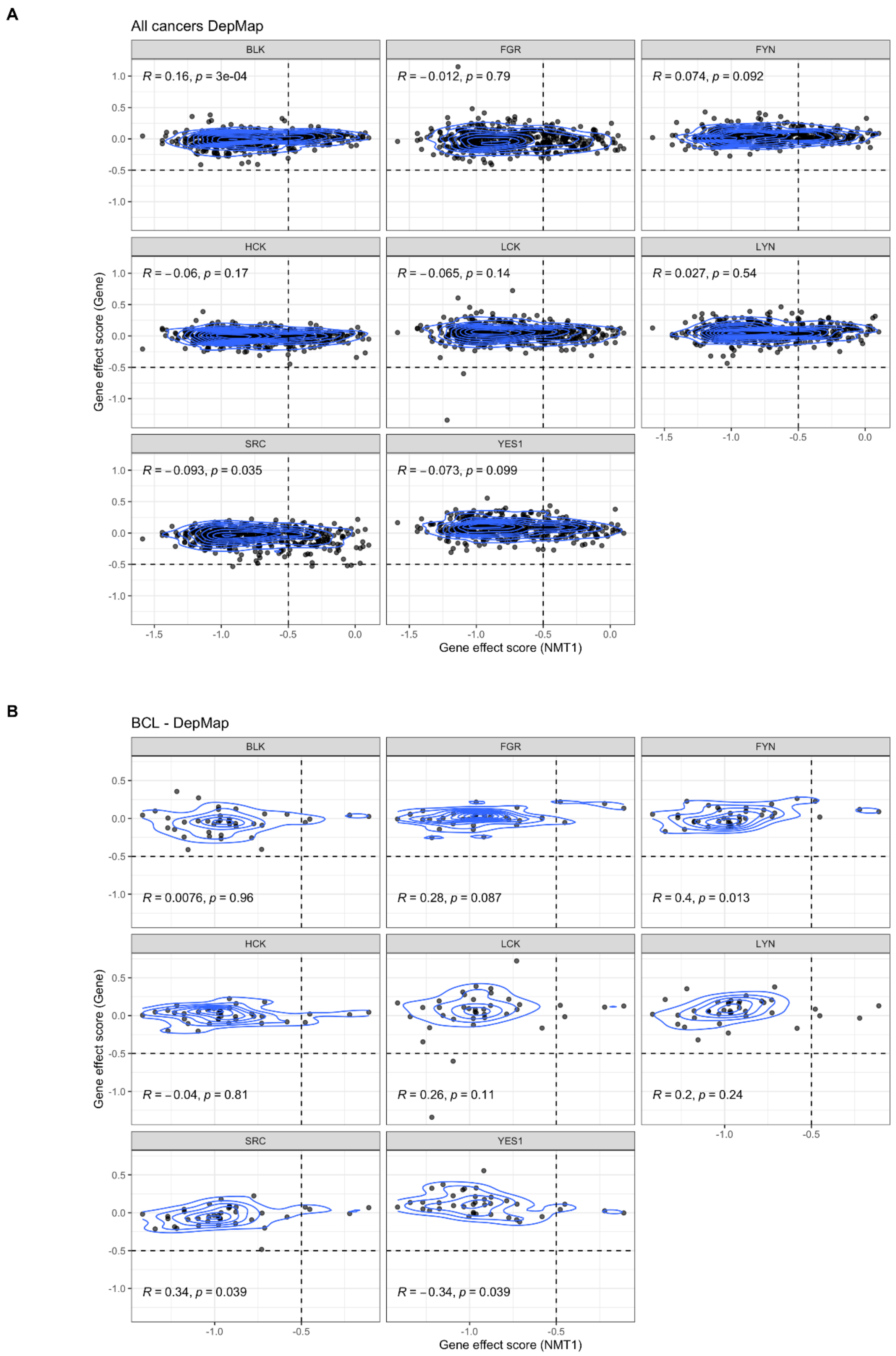

**Supplementary Figure 22: Correlation of gene effect scores between NMT1 and SFKs.**

**(A)** Pan-cancer correlation of gene effect scores between NMT1 knockout and the knockout of SFKs (Spearman rank test). **(B)** Leukemia and lymphoma correlation as in (A).

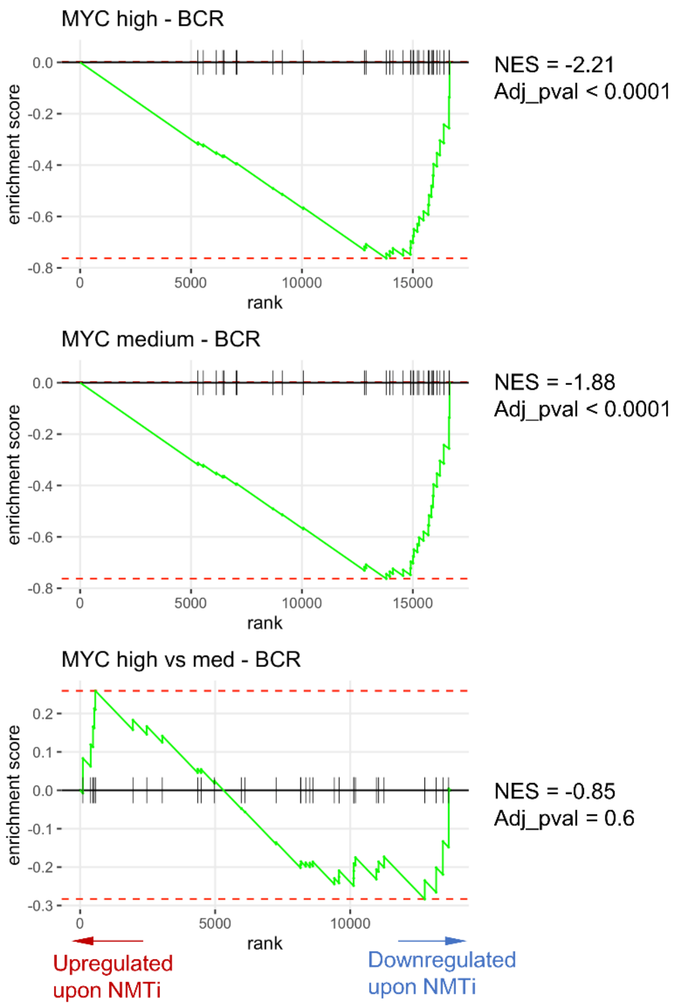

**Supplementary Figure 23: GSEA for BCR gene set in P493-6 cells treated with NMT inhibitor.**

GSEA for BCR gene set in P493-6 cells treated for 24 hours with NMT inhibitor IMP1088 (100 nM) with high (top), medium (middle) and high versus medium (bottom) MYC levels.

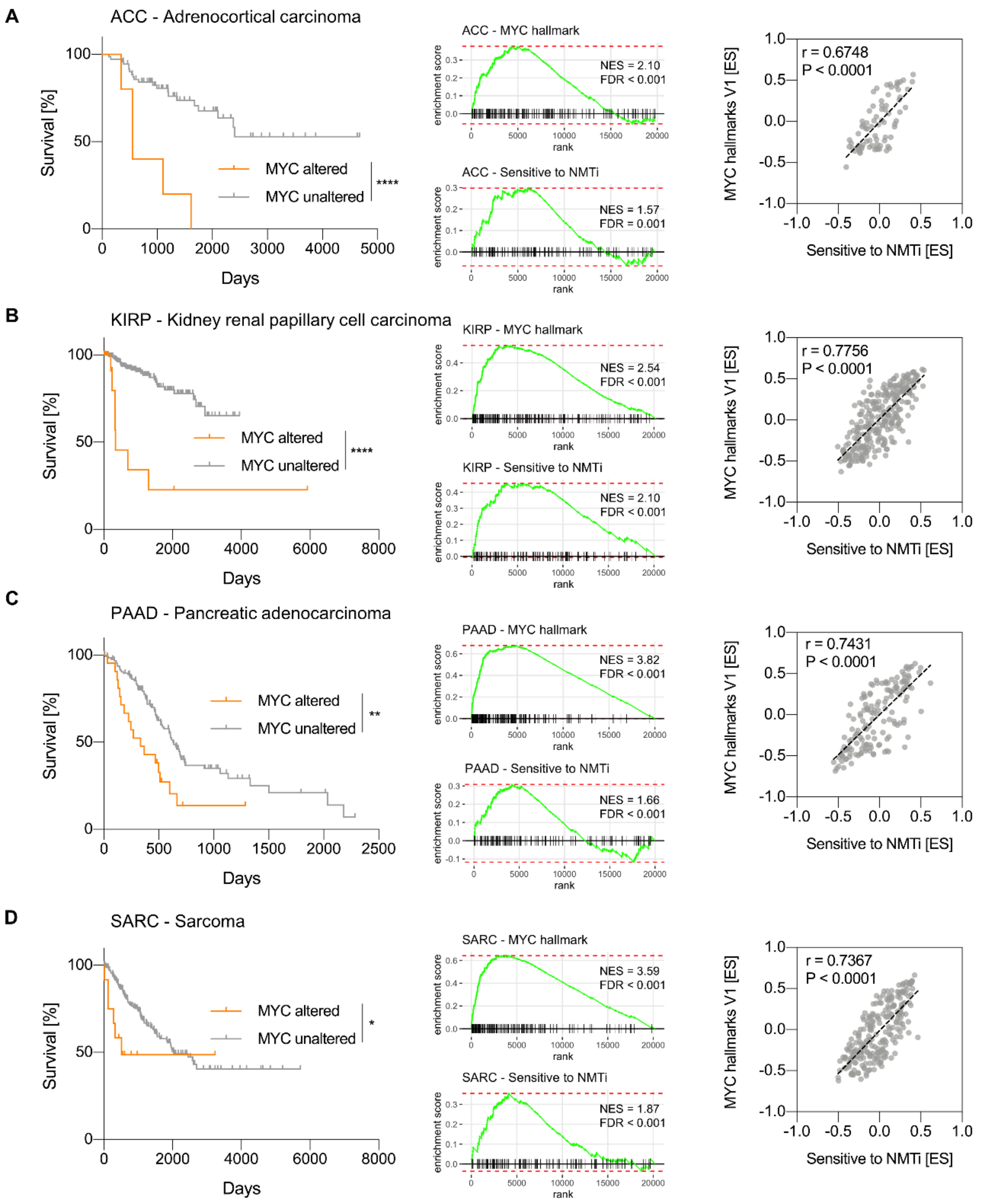

**Supplementary Figure 24: Presence of alterations in the MYC pathway correlate with worse clinical outcome in TCGA cohorts and with the "Sensitive to NMTi" gene set.**

**(A)** Left: correlation between the presence of alterations in the MYC pathway (extracted from *(15)*) with clinical outcome (extracted from *(14)*) in the TCGA AGG (Adrenocortical carcinoma) cohort (Mantel-Cox test). Middle: enrichment in patients with alterations in the MYC pathway for the "Hallmark MYC" gene set (top) and the "Sensitive to NMTi" gene set. Right: correlation between the ES of the "Hallmark MYC" gene set and "Sensitive to NMTi" gene set are shown, as determined by GSVA (Spearman rank correlation). **(B)** to **(D)** As in (A); **(B)** for the TCGA KIRP (Kidney renal papillary cell carcinoma) cohort; **(C)** for the TCGA PAAD (Pancreatic adenocarcinoma) cohort; and **(D)** for TCGA SARC (Sarcoma) cohort.

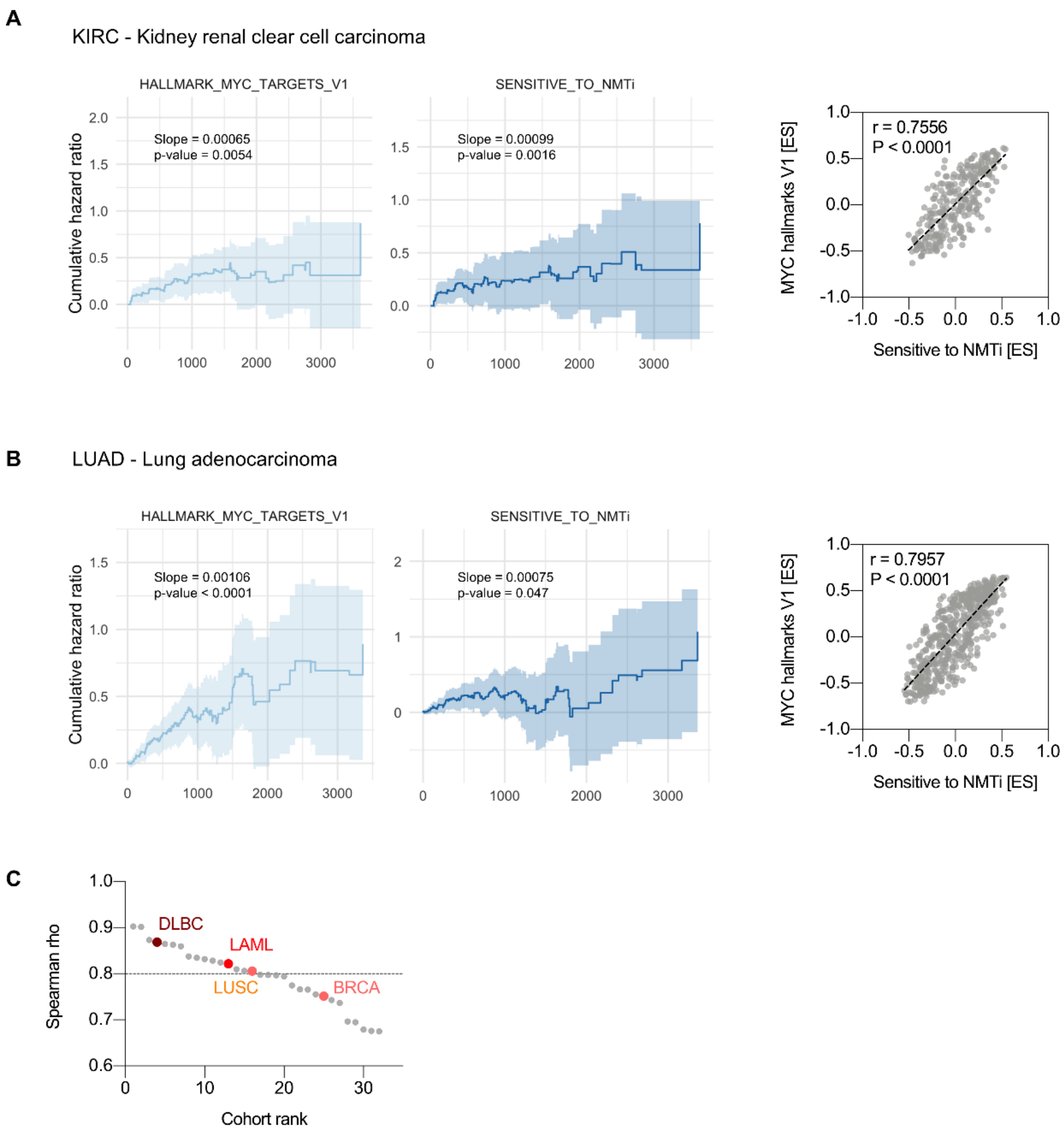

**Supplementary Figure 25: Expression of the "Hallmark MYC" gene set or the "Sensitive to NMTi" gene set correlate with worse clinical outcome for the TCGA cohorts KIRP and LUSC.**

**(A)** Left: correlation between the expression of the "MYC Hallmark" gene set, measured as GSVA ES, and clinical outcome for the TCGA KIRC (Kidney renal clear cell carcinoma) cohort. Middle: correlation between the expression of the "MYC Hallmark" gene set, measured as GSVA ES, with clinical outcome (Nelson-Aalen estimator). Right: correlation between the ES of the "Hallmark MYC" gene set and "Sensitive to NMTi" gene is shown (Spearman rank correlation). **(B)** Left: correlation between the expression of the "MYC Hallmark" gene set, measured as GSVA ES, and clinical outcome for the TCGA LUAD (Lung adenocarcinoma) cohort. Middle: correlation between the expression of the "MYC Hallmark" gene set, measured as GSVA ES, with clinical outcome (Nelson-Aalen estimator). Right: correlation between the ES of the "Hallmark MYC" gene set and "Sensitive to NMTi" gene are shown (Spearman rank correlation). **(C)** Spearman rank correlations between the ES of the "MYC Hallmark" and "Sensitive to NMTi" gene sets are shown for the tested TCGA cohorts, with a median correlation of 0.8 (dotted line) across all TCGA cohorts, and selected cohorts highlighted.

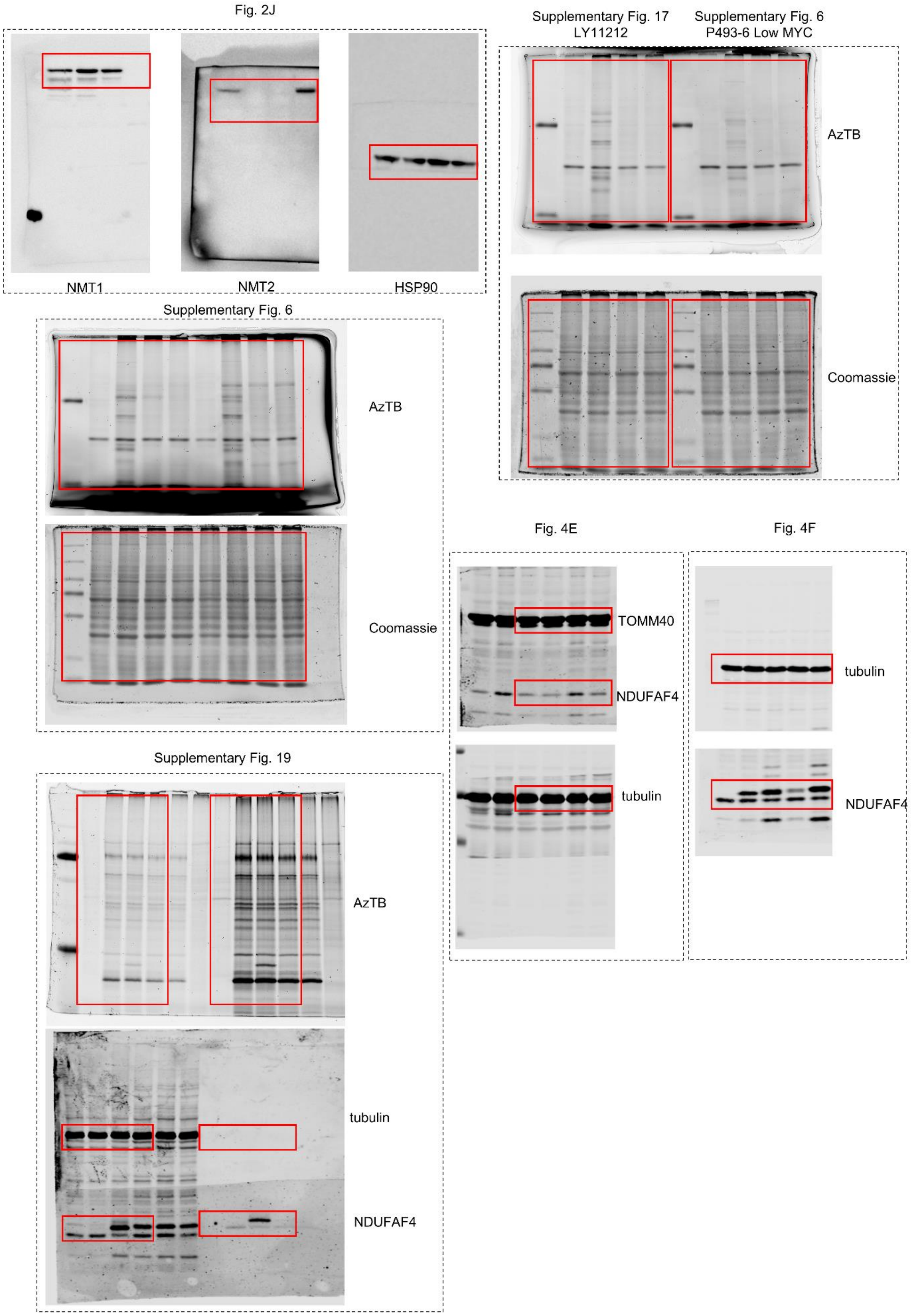

**Supplementary Figure 26: Uncropped western blots and gels.**

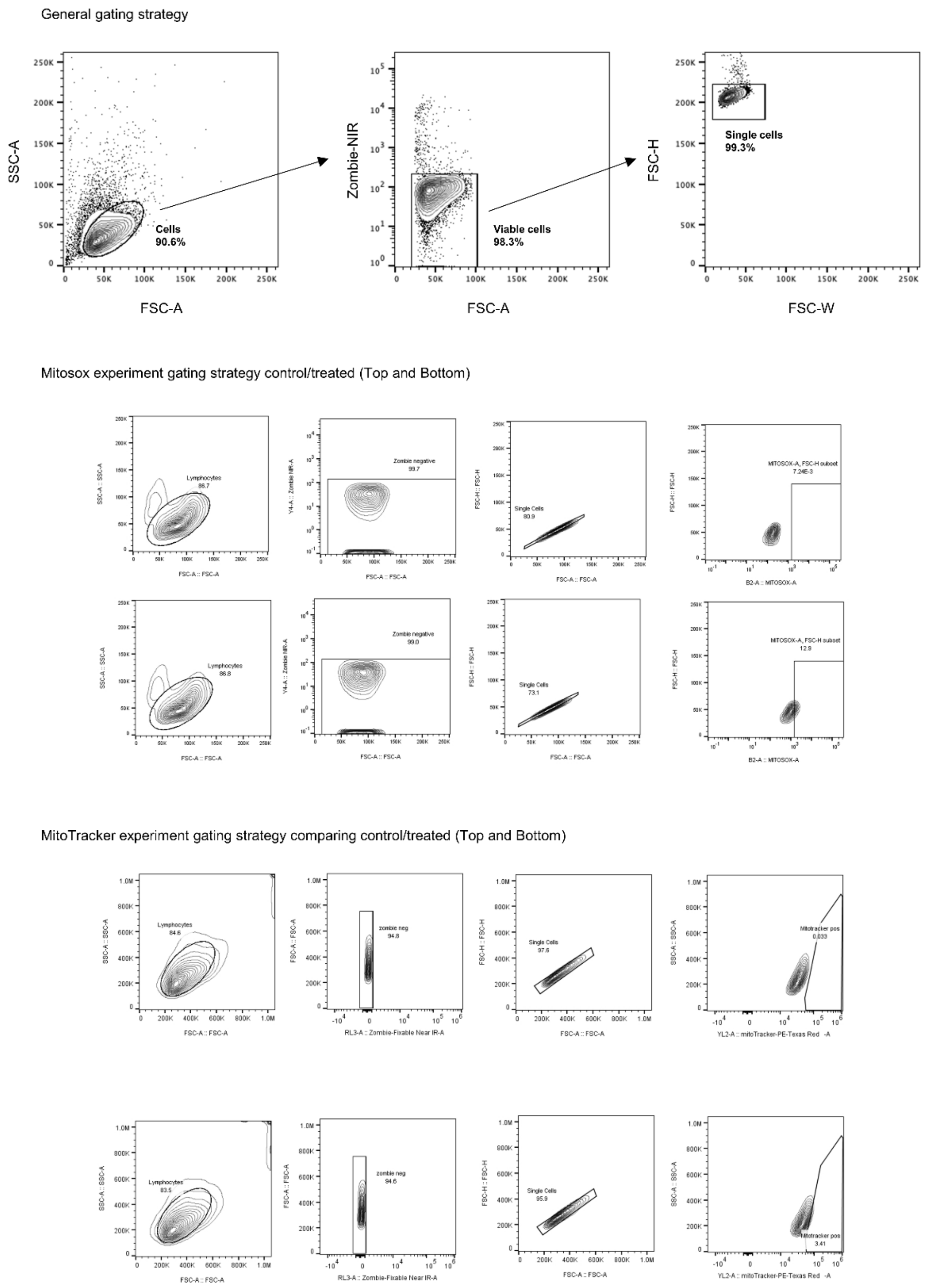

**Supplementary Figure 27: Examples of gating strategies for flow cytometry experiments.**

#### Supplementary methods

##### Enzymatic NMT assays

Full-length human NMT1 and NMT2 were produced as described previously *(3)* and used at the final concentration of 300 ng/mL *(4)*. The assay buffer (pH 7.8) contained 5 mM phosphate buffer, 0.125 mM EDTA, 0.025% Triton X-100 and 1% DMSO. The assays were performed in 96-well black plates (110 μl total volume) using the following dual NMT inhibitors: DDD86481, IMP1031, IMP1036 (synthesis reported in *(5)*) and IMP1088. Reactions were run for 30 min at room temperature and quenched with 60 μl 100 mM acetate buffer pH 4.8. NMT activity was determined through fluorescent detection of CoA-SH with 8 μM CPM ($\lambda_{ex}$ 380 nm, $\lambda_{em}$ 470 nm, EnVision 2102 multilabel plate reader, PerkinElmer), formed during the *N*-myristoylation of 4 μM SRC peptide (amino acids 2-16), using 4 μM myristoyl-CoA *(6)*. After background correction, $IC_{50}$ values were determined by fitting the four-parametric variable slope function in Prism (GraphPad). $IC_{50}$ values of DDD85646 for NMT1 and NMT2 are reported elsewhere *(7)*. For NDUFAF4 peptide (amino acids 2-10) assays the concentrations of WT and A3P peptides were varied. After background correction, Km values were determined by Michaelis-Menten function (nonlinear regression fit) in Prism (GraphPad). All assays were performed in three independent experiments.

##### Surface plasmon resonance (SPR)

Kinetics data were collected on Biacore S200 Biosensors (Cytiva). Recombinant human *N*-myristoyl transferase 1 (NMT1) 109-496aa proteins were immobilized to Series S CM5 sensorchips, which were maintained with a continuous flow of HBS-P+ (10 mM 4-(2-hydroxyethyl)-1-piperazineethanesulfonic acid adjusted to pH 7.4, 150mM NaCl and 0.5% (v/v) polysorbate 20). 7 min, 10 μL/min injections of 0.5 M 1-ethyl-3-(3-(dimethylamino)propyl)-carbodiimide (EDC) and 0.1M N-hydroxysuccinimide (NHS) facilitated catalytic activation of the carboxylate moieties of the surface dextran matrix. Recombinant NMT1 was exchanged into 10 mM Sodium Acetate pH5 buffer, at a final concentration of 10 μg/mL. Proteins were injected across the activated dextran surface for 50 s at 10 μL/min to achieve a low density immobilization (500-1000 RU). Unreacted succinimide esters were deactivated with a 7 min injection of 1 M ethanolamine at 10 μL/min. For each protein-coupled flow cell, a reference flow cell was also prepared. Compounds were stored at 50 mM in 100% DMSO at 25 °C. Stocks were diluted to 100 μM in DMSO, before being diluted to 1 μM + 1% DMSO in HBS-P+. Compounds were diluted further to 10 nM, followed by a 5-point, 2-fold serial dilution (10 nM, 5 nM, 2.5 nM, 1.25 nM 0.625 nM) in HBS-P+ + 4μM Myristoyl Coenzyme A + 1% DMSO (“Assay Buffer”). Solvent correction samples were made in HBS-P+ with 4 μM myristoyl-coenzyme A with an 8-point increasing concentration of DMSO from 0.05% to 1.5%. The diluted compound series was injected in order of ascending concentration for 400 s, across a specific flow cell, separated by a 50 s flow of assay buffer. Dissociation was monitored during a 1500 s injection of assay buffer. An equivalent series of injections was carried out using Assay Buffer, which was used to correct flow cell-specific artefacts.

**Public data sets**

The pre-processed microarray data, coding variants and copy number alterations for analysis of data from the Sanger institute were obtained from the GDSC project *(8)*. The RNA-seq data for the analysis of data from the Broad institute were obtained from the Cancer Cell Line Encyclopaedia *(9)*; Release 4Q2018). The CRISPR data for gene essentiality was obtained from the Broad DepMap project (*(10)*; Release 4Q2018) and from the Sanger Project Score project *(11)*. Gene sets used in this study were obtained from the MSigDB, namely the Hallmark gene sets and canonical pathways (*(12,13)*). IMP366 NMTi sensitivity data is available on the GDSC portal (https://www.cancerrxgene.org/) under the name ICL1100013. The RNA-seq for the TCGA analysis was obtained from the TCGA Research network by the NIH (https://gdc.cancer.gov/about-data/publications/pancanatlas ). The clinical outcome data for the TCGA was extracted from the following publication *(14)* and the information if the MYC network is altered from *(15)*.

**Gene set enrichment analysis (GSEA)**

After dividing the samples into two groups (sensitive or resistant to a given NMT inhibitor; highly or lowly dependent on NMT1; etc.) the signal-to-noise ratio was used to create a pre-ranked list of genes, according to the initial GSEA paper *(16)*. GSEA was subsequently applied and visualized with the R library fgsea *(17)*. Settings were kept to a standard: minSize = 15, maxSize = 500, nperm = 10,000. The 'Sensitive to NMTi' gene set was derived as the intersection of the leading-edge genes of the 10 gene sets with highest positive NES (most enriched in the sensitive cell lines). The CytoScape visualization of the gene sets positively enriched in the sensitive cancer cell lines was done according to a published protocol *(18)* after rerunning the GSEA on the published JavaScript-based program (GSEA version 3.0), with EnrichmentMap *(19)* and CytoScape *(20)*. The settings were: FDR cut-off = 0.1, Jaccard Overlap = 0.375, k-constant = 0.5.

**Gene set variation analysis (GSVA)**

The R library GSVA *(21)* was applied to estimate the enrichment score (ES) of selected gene sets for each cancer cell line in the Sanger microarray data, and in the TCGA RNA-seq data set. For the microarray data, the 50% most lowly variant and lowly expressed genes were removed by calculating the inter-quantile range (IQR) for each gene and removing the genes with the lowest IQR. For the RNA-seq lowly abundant transcripts were removed based on the threshold $\log2(x+1) \leq 0.5$. For both RNA-seq and microarray a "Gaussian" kernel was applied, and the enrichment score (ES) is reported as magnitude difference between the largest positive and negative deviations of the random walk. The GSVA was run, with the following settings: method = "gsva", kcdf = "Gaussian", mx.diff = TRUE, min.sz = 15, max.sz = 500.

**Identification of genes correlated to $IC_{50}$**

We identified genes correlated to compound IC50 by using Statistical Analysis for Microarrays (SAM), 'quantitative SAM' as implemented in the R statistical package samr (https://statweb.stanford.edu/~tibs/SAM). Quantitative SAM offers the possibility to identify genes whose expression profile correlate with a continuum if IC50 either negatively or positively. Genes were considered significant at an FDR≤10%.

**IPA Analysis**

In order to test whether genes correlated to $IC_{50}$ were representing biologically relevant functional trends we used a combination of the web-based tool DAVID and the Ingenuity Pathway Analysis (IPA) software *(22,23)*. More specifically, genes positively (or negatively) correlated with $IC_{50}$ were used as an input of the DAVID tool (Functional Clustering) to identify gene ontology terms enriched in the input gene lists. Clusters with enrichment score >1.5 were selected and representative gene ontology terms with FDR<10% within each cluster.

Factors that are likely to mimic the expression signatures linked to drug sensitivity were identified by using the IPA Upstream Regulator Analysis. In this analysis the software compares prior knowledge of expected effects between transcriptional regulators (e.g., transcription factors, growth factors, metabolites, drugs, etc.) and their targets with the transcriptional changes identified in a given analysis. In this analysis the software computes two measures, the p-value of overlap and the activation score. This approach has been used in our analysis to identify putative factors predicted to be active in the presence of cell lines that are highly sensitivity to the drug as well as in cell lines that are not very sensitive. Factors with an activation score >|3| and with a significant overlap *P*-value ($p<10^{-2}$) were selected for interpretation.

**Testing effects of presence of mutations in NMT substrates**

The $EC_{50}$ values of the NMT inhibitors **1** and **2** was first log10 transformed and then standardized to yield normally distributed potency values. This was followed by testing for each NMT substrate (n = 125) if the presence of a mutation had a significant effect on the potency between the mutated and not-mutated cancer cell lines with an ANOVA test. The *P*-values were corrected for multiple hypothesis testing with a Benjamini-Hochberg procedure.

**Cell culture**

All cells were cultured in a humidified 37 °C incubator at 5% (v/v) $CO_2$ atmosphere. DMEM medium (with GlutaMAX) was supplemented with 10% FCS, 10 mM HEPES, 1 mM sodium pyruvate, penicillin-streptomycin (100 U/ml and 100 µg/ml, respectively), 0.1 mM non-essential amino acids and 25 µM β-mercaptoethanol.

**Patient derived cells**

LY11212 was derived from a patient diagnosed with DLBCL, classified as GCB using nanostring, $CD20^+$, $CD10^+$, $BCL6^+$, $IRF4^{neg}$, $BCL2^+$, $MYC^+$, Ki67 80% by histology and $CD19^+$, $IgM^+$, $lambda^+$ by flow cytometry, with BCL2 translocation t(14;18)(q32;q21) and MYC translocation t(8;14)(q24;q32) by FISH.

**Cell viability measurements**

CellTiter-Blue (Promega G8080) was employed to assess cell viability following exposure of the cells to different concentration of the respective NMT inhibitors. The experiments were carried out in 96-well flat bottom plates with three or four technical replicates. P493-6 cells were cultured for 24 h in the respective MYC conditions (high-MYC: DMEM, low-MYC: DMEM with 0.1 µg/mL of doxycycline, medium-MYC: DMEM with 0.1 µg/mL of doxycycline and 1 µM of β-estradiol) at concentration of $5{\cdot}10^5$ cells/ml. The following day, 50 µl of cells was added to 50 µl of culture medium containing twice the concentration of inhibitors or control and the respective medium-MYC conditions. Shep-ER-MYCN cells were seeded at concentration of 1,000 cells/mL in 50 µl of medium (DMEM) and left to adhere overnight. The following day, 50 µl of culture medium containing 200 nM of 4-OH-Tamoxifen was added to the cells for 24 h. 100 µl of culture medium containing 100 nM 4-OH-Tamoxifen and twice the concentration of NMTi or controls were added to the respective wells. PDX LY11212 cells were seeded at $5{\cdot}10^5$ cells/ml with 100 µl of culture medium containing the range of inhibitor concentrations or DMSO control. HeLa cells were seeded in 50 µl at 1,700 and 2,000 cells/well respectively in quadruplicate on 96 wells flat bottom. The following day, 50 µl with twice the concentration of

IMP1088 were added to the cells and 72 h later CellTiter-Blue was added following the manufacturer's instructions. Once the experiment reached the endpoint, 10 or 20 μl of CellTiter-Blue was added to each well and the plates were left at 37 °C in the incubator for 2 h prior to measuring fluorescence at excitation/emission $\lambda_{ex}$ 560 nm, $\lambda_{em}$ 590 nm using the EnVision 2102 multilabel plate reader (PerkinElmer). Cell viability was normalized against the positive control (mixture of staurosporin and puromycin at the final concentration of 1 µg/mL and 10 µg/mL respectively). Prism (GraphPad) was used to fit the four-parametric variable slope function. Every experiment was repeated three times.

**Flowcytometric Analysis**

Single cell suspension cells were seeded in 96 U-well plates at the same concentration and conditions described for CellTiter-Blue experiments. For time course beyond 24 h, 100 μl of culture medium containing the respective MYC conditions, inhibitors, and control, were added to the cells. Shep-ER-MYCN were seeded at 7,000 cells/mL in 24 well plates flat bottom. 24 h later, culture media containing 200 nM 4-OH-Tamoxifen or EtOH as control were added to the wells for 24 h. Culture medium was removed, and fresh medium containing 100 nM 4-OH-Tamoxifen, NMTi and DMSO controls were added to the respective wells for the time course indicated. Edu (Thermo Fisher Scientific, F10347) was added to the cells 2 h before fixation at the final concentration of 10 μM.

For all the cells, caspase activation was monitored with α-active caspase-3 antibody (C92-605, 550821, BD Bioscience), DNA content was stained with FxCycle Violet (Thermo Fisher scientific, F10347), and proliferation was visualized employing Click-iT EdU AlexaFluor 488 Flow Cytometry Assay kit. To exclude dead cells, Zombie NIR (423105, Biolegend) was used according to the manufacturer instructions. The cells were fixed with 4% PFA, and permeabilized employing CytoFix/Cytoperm (BD Bioscience 554714) and washed in perm/Wash Buffer (BD Bioscience, 554723). MACSQuant VYB (Miltenyi Biotech) was used to acquire the samples and analyzed using FlowJo software.

#### Cloning

gRNA_cloning vector was a gift from George Church (Addgene plasmid 41824) *(24)*, and pCas9_GFP was a gift from Kiran Musunuru (Addgene plasmid 44719) gRNA were designed using ChopChop tools *(25,26)* and synthetized according to the Church lab protocol.

Full-length human NDUFAF4 gene was ordered as a 525 bp geneblock from Integrated DNA Technologies. The NDUFAF4 gene was cloned into a C-FLAG pcDNA3 vector by restrictionless cloning using KOD polymerase for expression as a C-terminal FLAG-tagged construct *(27)*. To incorporate the G2A and A3P mutations, corresponding substitutions were introduced into forward primers used for restrictionless cloning. The inserts were confirmed by DNA sequencing. Primers used for cloning and sequencing are listed in Supplementary Table 2.

#### Generation of NMT1 and NMT2 CRISPR-Cas9 knockout clones in HeLa cells

HeLa cells were obtained from ATCC and verified by STR at the Francis Crick Institute Cell Services. Cells were cultured in a humidified 37 °C incubator at 5% (v/v) $CO_2$ atmosphere in DMEM with low glucose and 10% (v/v) FCS, supplemented with HEPES, sodium pyruvate, penicillin-streptomycin, and non-essential amino acids. $5 \cdot 10^6$ cells/mL were seeded on 10 cm dishes. 24 later, 2 μg of pCas9_GFP plasmid alone (control) or in parallel with 2 μg of the respective gRNA (Supplementary Table 2) was transfected using Lipofectamine 2000 (Thermo Fisher Scientific 11668019) according to the manufacturer's instructions. 48 h later, cells were harvested and sorted by BD FACSAria III according to the GFP fluorescence at Imperial College South Kensington facility, in 96 flat-bottom well plates to generate single cell clones. Clones where lack of NMT1 and NMT2 protein was verified by western blot, were sequenced to confirm successful knockouts.

#### Verification of CRISPR-Cas9 knockouts by sequencing

Genomic DNA (gDNA) was extracted from every clone using DNA extraction kit (QIAGEN 69504) according to the manufacturer's instructions. gDNA concentration was measured using a NanoDrop 2000c spectrophotometer (Thermo Scientific). Because the final PCR product was used for sequencing analysis, PCR amplification reactions were performed using Phusion High Fidelity Polymerase (New England BioLabs M0530). Primers sequence are indicated in Supplementary Table 2. PCR was performed in a final volume of 25 µL, containing 10 µL of 2x PCR Master Mix, 1.25 µL of forward primers, 1.25 µL of reverse primers, 40 ng of gDNA, and water. PCR amplifications were performed according to the following parameters: 98 °C for 30 s; 25–35 cycles (suitable cycles were chosen for each gene) of 98 °C for 5-10 s, 56–65 °C (proper annealing temperature was chosen for gene) for 10 s, and 72 °C for 15-30 s, with a final extension step of 72 °C for 5 min. The PCR product was then purified and sent for sequencing to Genewiz (Sanger Sequencing). Chromatograms were analyzed using FinchTV.

**NDUFAF4 biochemistry**

HEK293 cells were plated in 6-well plates and transfected with NDUFAF4-FLAG constructs (WT, G2A and A3P) using 2 µg of DNA plasmid and 6 µl of Fugene HD transfection reagent (1:3 ratio) per well. 24 h after transfection, cells were treated with 10 µM of MG132 or DMSO as a vehicle control for a further 16 h. Cells were lysed (lysis buffer: PBS with 1% Triton X-100, 0.1% SDS and 1x Complete EDTA-free protease inhibitor cocktail (Roche)) and subjected to western blotting.

**YnMyr labelling for in-gel fluorescence and streptavidin pulldown**

P493-6 (low-, medium-, high-MYC) and LY11212 cells were treated with 100 nM IMP1088, 1 µM DDD86481 or DMSO (negative control) for 30 min, followed by YnMyr (20 µM) and incubated for 18 h. HEK293 cells were treated with YnMyr (20 µM) for 18 h. Lysates (same lysis buffer as above) were subjected to ligation with 0.1 mM Azido-TAMRA-Biotin (AzTB) in a click reaction buffer containing 1 mM $CuSO_4$, 1 mM TCEP for 1 h and analyzed by in-gel fluorescence using the Typhoon FLA 9500 imager (GE Healthcare). For streptavidin pulldown, cell lysate protein was chloroform-methanol precipitated, resuspended in 0.1% SDS and 5 mM DTT and Streptavidin MyOne beads (Thermo Scientific) were added for 2 h (1000 RPM shaking at room temperature). Consistent gel loading was confirmed by Coomassie stain

(Instant Blue, Expedeon) or by western blot using tubulin as a loading control, scanned using Odyssey CLx imager (Li-COR) and analyzed by ImageStudio software (Li-COR).

**Western blotting**

Protein samples were prepared with 4x Laemmli sample loading buffer (BioRad 1610747) and 10% β-mercaptoethanol, boiled for 5 min at 95 °C and resolved on 10% or 12% (w/v) SDS-PAGE gels running at 180 V. For NMT KO experiments, proteins were transferred onto nitrocellulose membrane for 1.5 h using wet blotting (BioRad) and blocked for 1 h with 5% (w/v) skimmed milk in TBS 0.1% Tween-20. Proteins were detected using NMT1 (Sigma Aldrich, HPA022963), NMT2 (BD 611310), and GAPDH (Abcam ab9485) was used as loading control. HRP-conjugated secondary antibodies (Advansta, R-05071-500 and R-05072-500) were used to detect proteins by ImageQuant LAS4000 (GE), quantified by Image J (NIH Bethesda), and normalized against the loading control. For NDUFAF4 experiments, proteins were transferred onto nitrocellulose membrane using Trans-Blot Turbo (BioRad) and blocked for 30 min with 5% (w/v) BSA in TBS 0.2% Tween-20. Proteins were detected using NDUFAF4 (ABclonal, A14345), TOMM40 (Proteintech, 18409-1-AP), α-tubulin (Sigma, T5168). Fluorescence signal from secondary antibodies (LI-COR) was quantified using the Odyssey LI-COR system. Quantification was performed using Image Studio software (LI-COR) and statistical analysis was performed using Prism (GraphPad).

**RNA-seq**

P493-6 cells were cultured overnight as previously described in the respective MYC conditions (high- and medium-MYC). After 24 h of induction of the respective MYC levels 100 nM IMP1088 or a respective DMSO control were added to the cells for 24 h, before further processing. The samples were produced in biological quadruplicate and lysed with the QIAzol lysis reagent. The RNA was extracted with the miRNeasy kit (Qiagen) according to manufacturer's instructions. The sequencing was performed in the Advanced Sequencing facility in the Francis Crick Institute. The sequencing was performed on an Illumina HighSeq system, the read length was paired-end 100 bp, and the sequencing depth was 25 to 30 million reads per sample. For the total RNA-seq library preparation the KAPA RNA HyperPrep Kit was used. Quality control of the reads and read trimming was performed with Trimmomatic (version 0.36) *(28)*. Alignment of the reads was carried out with STAR (version 2.5.2a) against the human transcriptome Ensembl GRch38 *(29)* (release 86); counts per gene and sample were obtained using RSEM (version 1.2.31) *(30)*. Gene level differential expression analysis was done using the program DESeq2 (version 1.18.1) *(31)*; genes with an adjusted *P*-value of <0.05 were considered statistically significant.

**Proteomics sample preparation**

P493-6 cells were cultured overnight as previously described in the respective MYC conditions (high- and medium-MYC) The following day, cells were centrifuged, washed twice in warm PBS, and divided into two flasks, containing 100 nM IMP1088 or the respective DMSO control in SILAC DMEM (Sigma) supplemented with the respective MYC conditions, heavy arginine and lysine (Cambridge Isotope Laboratories) 10% dialyzed FBS (Sigma), and 1% penicillin-streptomycin. Cells were collected at intervals of 4 h, centrifuged, washed in PBS and the pellet snap frozen until further analysis. For the spike-in SILAC, P493-6 were grown in R6K4 DMEM medium (Cambridge Isotope laboratories) supplemented with 10% dialyzed FBS and 1% penicillin/streptomycin for 6 passages. The incorporation of the R6K4 label was determined to be 95%. Cells were then cultured overnight in the respective MYC conditions, pulled, and the pellet was washes and snap frozen in liquid nitrogen.

The pellet was dissolved in HEPES 50 mM pH 8, containing 0.1% SDS (w/v) and EDTA free protease inhibitor tablets (Roche 23225), cold sonicated two times for five seconds using a probe sonicator. Proteins were precipitated (MeOH:$CHCl_3$ at 4:1 ratio), the pellet was washed with cold 10% Water:MeOH, air-dried and resuspended in 50 mM HEPES pH 8. Protein concentration was determined with BCA protein kit (Thermo Fisher Scientific 23225). Lysates (50 μg) were combined in 3:1 ratio with R6K4 labelled cells lysates, digested and incubated overnight with a mixture of 0.1 μg Trypsin (Promega V511A), 5 mM TCEP and 10 mM CAA at 37 °C with gentle shaking. The following day, the reaction was quenched with 0.1% TFA (w/v), and samples were fractionated according to the protocol described in *(1)*, and analyzed by nanoLC-MS/MS on a Thermo Q-Exactive instrument as described previously *(1,4)*.

**Proteomics initial data analysis**

The data were processed using MaxQuant (version 1.5.6.5), using the inbuilt Andromeda search engine. The MS/MS spectra were matched against the human reference proteome with isoforms (UniProt, accessed July 2016). Cysteine carbamidomethylation was defined as a fixed modification; methionine oxidation and N-terminal acylation were set as variable modifications. As digestion mode 'Trypsin/P' was chosen, and a maximum of two missed cleavages allowed. Both the options 'match between runs' and 'unique and razor peptides' for protein quantifications were selected. The processed data was then further analyzed with Perseus (version 1.5.6.0). For all experiments the protein groups that are only identified by one side, that are potential contaminants, or that matched to the reverse database were excluded from further analysis. Per time point and condition (e.g., high-MYC cells treated with DMSO for 24 h) protein groups were removed only if a single SILAC ratio (be it H/L, H/M or M/L) was identified, i.e., only a value for a single time point obtained. Further analysis was conducted in R.

**Proteomics - imputations**

To impute missing data points (e.g., in the case of having three out of four time points), different machine-learning regression learners were tested, using the mlr library in R, based on trees and/or linear regression models (regr.rpart, regr.xgboost, regr.gbm, and regr.cubist), and test different ways of multiple imputation (a cross-validation like approach, bootstrapping and subsampling). The hyperparameters for regr.xgboost were set to: nrounds = 10, eta = 0.3, max_depth = 10, subsample = 0.5. Multiple imputations were applied to reflect to some extent the uncertainty of the underlying data. Firstly, the data sets were split into protein groups with or without missing values (NAs). Secondly, different sampling methods were applied to these two resulting data sets with or without NAs, and subsequently combined again. This way a situation was avoided in which for a given imputation iteration the number of data points with or without NAs did not represent the distribution of actual missing values of the original data. For the cross-validation like approach the data set was initially randomized, then cut into 10 equal pieces. For the multiple imputation one piece after the next was withheld from the imputation. For the bootstrapping, over 50 iterations were bootstrapped, and imputed for each bootstrap. For the resampling, random 80% of the original data was resampled over 50 iterations and impute on each resulting subsample. To assess how well the imputed values out of the different learners and different multiple imputation methods reflect the none-imputed data, the distance metric of the Kolmogorov-Smirnov test was used. The higher this distance metric between the data points of the none-imputed vs. the imputed values, the less do the imputed values match the reference distribution of the none-imputed values, indicating worse imputation. This metric was used to select for the learner 'regr.cubist' with a 50-fold bootstrapping as a standard imputation which was used in subsequent analysis.

**Proteomics – dynamics in synthesis or degradation**

To calculate for a given time point the difference in proteome dynamics, either synthesis (described by the H/M ratio) or degradation (described by the M/L ratio) a similar strategy as used for multiplexed proteomics for protein dynamics was applied *(32)*. Savitski et. al. used the number of quantified TMT spectra to bin their data by data quality. In this study, as triplex-SILAC was applied, the geometric mean of the 'quantified spectra + 1' for each data point in a given comparison was used for the binning strategy. (We added +1 to account for cases in which the missing values were imputed via the described strategy – see above.) The geometric mean has the advantage of being outlier-insensitive, thus, a case in which only a single time point for a single replicate has a larger number of quantified spectra would still be put into low data quality bin. Based on the distribution of geometric means observed in the data, 5 cut-offs were chosen for the geometric mean: 1, 4, 8, 16, 32. These cut-offs also ensured a minimum of 250 samples per bin.

Subsequently, the approach from Savitski et. al., was applied. In brief, for each bin the standard deviation from the distribution of differences between the difference of $\log_2$-

transformed fold changes for the same protein divided by the square root of two (see formula **1**) was calculated using robust estimation (based on the interquartile range, see formula **2** and *(33)*).

$$\frac{log_2(FC_{rep1}) - log_2(FC_{rep2})}{\sqrt{2}}$$

Formula **1**

$$SD \approx \frac{IQR}{2 \times \phi^{-1} \times \frac{0.75n - 0.125}{n + 0.25}} \approx \frac{IQR}{1.35}\ (for\ n > 150)$$

Formula **2**

(IQR = interquartile range; n = number of samples in the bin)

In each bin for each $log_2$-fold change a *P*-value was calculated based on a Z-test, using the previously robustly calculated standard deviation. Subsequently, the bins were combined, and multiple hypothesis correction (using the Benjamini-Hochberg procedure) was applied on the whole data set. A protein was considered significantly altered in its synthesis or degradation if, firstly, the adjusted *P*-values in both replicates was ≤ 0.05; secondly, if the change was in the same direction in both replicates; and thirdly, if the $log_2$ fold change greater than 0.37 (a 1.3-fold change in either direction). In the case, in which the high-MYC were compared to the medium-MYC cells, we additionally accounted for differential proliferation rates over time, and demanded higher log2 fold changes for the different time-points, based on the relative differences in proliferation rates (at 4 h - 0.4; at 8 h - 0.41; at 16 h - 0.45; at 24 h - 0.48)

**Proteomics – half-life calculations**

Protein half-lives were calculated based on work from other labs *(34)*. kA of a given protein group was calculated according to formula **3** and the half-life with formula **4**.

$$k_A = \frac{\sum_{i=1}^{t_n} \log(r_i + 1) \times t_i}{\sum_{i=1}^{t_n} t_i^2} - \frac{\ln(2)}{t_{dt}}$$

Formula **3**

($t_n$ = no of time points; $r_i$ = H/L ratio at time point i; $t_i$ = time point i; $t_{dt}$ = doubling time of the cell line)

$$T_{1/2} = \frac{\ln(2)}{k_A}$$

Formula **4**

For each protein group a linear model was fitted between log($r_i$+1) and the time points. Protein groups with coefficients of determination ($R^2$) of <0.9 were discarded. The quality-value was set to 'weak' if it was possible to determine a H/L ratio in three out of four time points; 'good' if the H/L ratio in three out of four time points were based on a minimum of three quantified peptides per H/L ratio; and 'poor' for everything else (as in *(34)* and *(35)*).

**1D and 2D enrichments**

1D and 2D enrichments (the former to test for pathway enrichment in the proteomics dynamics solely, the latter to compare the protein synthesis dynamics with the abundance changes of RNA transcripts) were conducted with according to published procedures *(36)*. Enrichment of GO terms, downloaded from the MSigDB was tested *(12)*. A minimum threshold of overlap between the measured genes and the respective GO terms of 10 was applied; the FDR threshold for the 1D enrichment was set to 2%; for the 2D enrichment was set the FDR threshold to 0.1%.

**Cytofluorimetry for MitoTracker staining**

MitoTracker red (M22425) was used according to the manufacturer's instructions. Briefly, P493-6 were cultured overnight in the different MYC conditions as described previously. The following day, cells were transferred to a 96 U-well plates and treated for 18 h with 100 nM IMP1088. Next, the plate was centrifuged, the culture medium was carefully removed, and the pellet was resuspended in a culture medium containing 200 nM of MitoTracker staining for 30 min at 37 °C. The cells were then washed with PBS containing Zombie NIR for 5 min to discriminate the ones with intact membrane, re-pelleted and resuspended into prewarmed FACS buffer (PBS with 2% w/v FCS) and analyzed using MACSQuant VYB analyzers (Miltenyi Biotec). To analyze superoxide production, MitoTracker Red CMXRos (M7512) was used at the final concentration of 5 μM for 15 min. Cells were then treated as described for M22425 and run using MACSQuant. FlowJo software was used to analyze the data and the experiments were run in biological triplicate and every time in technical replicate.

**Oxygen consumption rate (OCR) measurements**

Oxygen consumption rate was measured using Seahorse XFe96 extracellular flux analyzer (Agilent Technologies) according to the manufacturer's protocols for suspension cells. $5 \cdot 10^5$ cells/ml of P493-6 cells were grown as described before in a 6-well plate in the different MYC conditions (medium- and high-MYC) for 24 h. $5 \cdot 10^5$ cells/ml LY11212 cells were grown as described before. NMT inhibitors (100 nM IMP1088 or 1 μM DDD86481) were added for an additional 12 or 18 h. Seahorse cartridge (Agilent Technologies) was hydrated in 200 μl XF calibrant solution (Agilent Technologies) overnight in a non-$CO_2$ incubator at 37 °C. On the day of the assay, the 96-well Seahorse cell culture plate (Agilent Technologies) was coated with 0.01% poly-L-ornithine solution (Sigma Aldrich) for 2 h prior to seeding $5 \cdot 10^4$ cells/well (P493-6) or $10^5$ cells/well (LY11212) in 50 μl of Seahorse XF base medium (unbuffered DMEM with phenol red), containing 10 mM glucose, 1 mM sodium pyruvate and 2 mM L-glutamine. To allow the cells to adhere to the bottom, the plate was centrifuged at 200g for 1 min with the lowest break setting. The seeded plate was placed into a non-$CO_2$ incubator at 37 °C for 20 min after which an additional 130 μl of the culture medium was added to the wells. The plate was then placed into a non-$CO_2$ incubator at 37 °C for 20 min before running the assay. The cartridge injection ports A, B and C were filled with 20 μl of oligomycin (15 μM), 22 μl of carbonyl cyanide-4 (trifluoromethoxy) phenylhydrazone (FCCP) (20 μM) and 25 μl of rotenone/antimycin A mixture (5 μM each) diluted in Seahorse XF base culture medium to give the final assay concentrations of 1.5 μM oligomycin, 2 μM FCCP and 0.5 μM rotenone/antimycin A (injected in the same order). Oligomycin, FCCP, antimycin A and rotenone were purchased from Sigma Aldrich. Mixing, waiting and measurement time were set to 2, 2 and 4 min respectively. The data were obtained using Wave software (Agilent Technologies) and analyzed with Prism (GraphPad).

**Xenografts**

Six-to-seven-week old female NSG (IL2R-NSG) were provided from the Francis Crick Institute BRF facility. $10^7$ cells/100 µl/body of LY11212 resuspended in 50% PBS and 50% Matrigel (Corning 356230)) were transplanted subcutaneously (s.c.) into the right flank of each female. After three days, NMTi was injected through intraperitoneal (IP) injection (final drug concentration 25 mg/kg in 200 µl) once a day. Tumor volumes were calculated as 1/2 × length × $width^2$. NMTi was resuspended in phosphate buffer containing 5% DMSO (Sigma D8418), 20% PEG400 (Hampton Research HR2-603) and 0.5% Tween-80 (Sigma P4780). For IMP1320 oral gavage experiment, SCID mice were inoculated subcutaneously in the right flank region with $5x10^6$ viable DoHH2 cells resuspended in 0.1 mL PBS mixed with Matrigel at 1:1 ratio. Mice were assigned to treatment groups when tumor volumes reached 100-150 $mm^3$ with ten mice per group. Oral QD dosing of Vehicle (10 mM $Na_2HPO4$ + 0.2% Tween-80) and 50 mg/kg IMP1320 commenced one day after of randomization, for ten consecutive days. Tumor volumes were measured twice weekly and calculated using the formula 0.5 ($LxW^2$). Statistical analyses used two-way ANOVA (PRISM, GraphPad). Animal experiments were carried out in accordance with national and institutional guidelines for animal care and were approved by The Francis Crick Institute biological resources facility strategic oversight committee (incorporating the Animal Welfare and Ethical Review Body) and by the Home Office, UK. All animal care and procedures followed guidelines of the UK Home Office according to the Animals (Scientific Procedures) Act 1986 and were approved by Biological Research Facility at the Francis Crick Institute.

**Survival analysis**

According to recommendation *(14)* each TCGA cohort was divided into a group of patients with and without alterations in the MYC pathway *(15)*, and tested the survival for the progression free interval (PFI) or overall survival (OS). The Cox-Hazard ratio between the two groups was determined with Prism (GraphPad), if there were at least five patients in each group with uncensored events. For the Nelson-Aalen estimator the R survival package (https://cran.r-project.org/web/packages/survival/citation.html) was applied and used as input the ES for the Hallmark MYC gene set or the ‘Sensitive to NMTi’ gene set in a given cohort, as determined by GSVA.

**Supplementary Table 2. gRNA, cloning and sequencing primers**

| Primer | Sequence |
|---|---|
| NMT1 gRNA 1F | TTTCTTGGCTTTATATATCTTGTGGAAAGGACGAAACACC**ggcgaagtggtgaacaccca** |
| NMT1 gRNA 1R | GACTAGCCTTATTTTAACTTGCTATTTCTAGCTCTAAAAC**tgggtgttcaccacttcgcc** |
| NMT2 gRNA 1F | TTTCTTGGCTTTATATATCTTGTGGAAAGGACGAAACACC **ggctgtgtacaccgcgggag** |
| NMT2 gRNA 1R | GACTAGCCTTATTTTAACTTGCTATTTCTAGCTCTAAAAC **ctcccgcggtgtacacagcc** |
| NMT2 gRNA 2F | TTTCTTGGCTTTATATATCTTGTGGAAAGGACGAAACACC**gaaaaactcaagtttggtat** |
| NMT 1F Sequencing | TCTTTGCCAGCAGAGAGGAT |
| NMT 1R Sequencing | CTGGCGGATATTGTCCTTGT |
| NMT2 1F Sequencing | ATAAGTGCCATCCCAGCAAA |
| NMT2 1R Sequencing | TGATCGATGCCAGTATCTGC |
| NMT2 2F Sequencing | GATGTATTCAATGCACTGGATT |
| NMT2 2R Sequencing | CACTTACCTTTTCAGAATCTGT |
| NDUFAF4 F WT | CTAGACTCGAGGGTACCGGATCCATGGGAGCACTAGTGATTCGCG |
| NDUFAF4 F G2A | CTAGACTCGAGGGTACCGGATCCATGGccGCACTAGTGATTCGCGGTATC |
| NDUFAF4 F A3P | CTAGACTCGAGGGTACCGGATCCATGGGAcCcCTAGTGATTCGCGGTATC |
| NDUFAF4 R | CTTGTCATCGTCGTCCTTGTAGTCTTTTGATCGTATTGCTTTCTTGTCTTCAGG |

NDUFAF4 geneblock:

atgggagcactagtgattcgcggtatcaggaatttcaacctagagaaccgagcggaacgggaaatcagcaagatgaagccctctgtcgctcccagacacccctctaccaacagcctcctgcgagagcagattagtctctatccagaagttaaaggagagattgctcgtaaagatgaaaagctgctgtcgtttctaaaagatgtgtatgttgattccaaagatcctgtgtcttccttgcaggtaaaagctgctgaaacatgtcaagagccgaaggaattcagattgccgaaagaccatcattttgatatgataaatattaagagcattcccaaaggcaaaatttccattgtagaagcattgacacttctcaataatcataaActtttcccagaaacctggactgctgagaaaataatgcaggaataccagttagaacagaaagatgtgaattctcttcttaaatattttgttacttttgaagtcgaaatcttccctcctgaagacaagaaagcaatacgatcaaaa
